# ClustSIGNAL identifies cell types and subtypes using an adaptive smoothing approach for scalable spatial omics clustering

**DOI:** 10.64898/2025.11.30.691081

**Authors:** Pratibha Panwar, Boyi Guo, Haowen Zhou, Stephanie C. Hicks, Shila Ghazanfar

**Affiliations:** School of Mathematics and Statistics, The University of Sydney, NSW 2006, Australia; Sydney Precision Data Science Centre, The University of Sydney, NSW 2006, Australia; Charles Perkins Centre, The University of Sydney, NSW 2006, Australia; Department of Biostatistics, Johns Hopkins Bloomberg School of Public Health, Baltimore, MD, USA; Division of Biostatistics, Department of Population Health Sciences, University of Utah, Salt Lake City, UT, USA; Bioinformatics and Systems Biology Graduate Program, University of California San Diego, La Jolla, CA, USA; Department of Biomedical Engineering, Johns Hopkins School of Medicine, Baltimore, MD, USA; Center for Computational Biology, Johns Hopkins University, Baltimore, MD, USA; Kavli Neuroscience Discovery Institute, Johns Hopkins University, Baltimore, MD, USA; Data Science and AI Institute, Johns Hopkins University, Baltimore, MD, USA

**Author notes:** Co-corresponding authors: Stephanie C. Hicks and Shila Ghazanfar.

**Keywords:** spatial clustering, unsupervised clustering, adaptive smoothing, data imputation, spatial transcriptomics

## Abstract

Unsupervised clustering of spatial transcriptomics data is challenging due to sparsity and complex tissue structures. We introduce ClustSIGNAL spatial clustering method that employs neighbourhood complexity-based expression smoothing. ClustSIGNAL uses entropy to measure neighbourhood compositions and generate cell-specific weights that control the effective smoothing radius within neighbourhoods. Simulation analysis demonstrates its robustness to segmentation errors and sparsity. Benchmarking across diverse SRT datasets shows its ability to perform scalable, multi-sample clustering of atlas-level data. By stabilising expression in homogeneous regions and preserving distinct expression in heterogeneous regions, ClustSIGNAL avoids over-smoothing and identifies biologically meaningful spatially-informed cell states. ClustSIGNAL Bioconductor package: bioconductor.org/packages/clustSIGNAL.

## Introduction

Technologies to profile the transcriptome have progressed from bulk to single-cell resolution, and now to spatially-resolved transcriptomics (SRT) where transcripts are captured *in situ* [1–6]. SRT technologies vary considerably depending on the combination of methodology, from imaging-based platforms that use gene panels (e.g., seqFISH [7], Vizgen MERFISH [8], 10x Xenium, Nanostring CosMx SMI [9]) to sequencing-based whole transcriptome capture approach (e.g., 10x Visium [10], Slide-seq [11], Slide-seqv2 [12]), and spatial resolution, from low-resolution spots containing multiple cells (e.g., 10x Visium, Nanostring GeoMx DSP [13]) to high-resolution subcellular information capture (e.g., 10x Visium HD [14], STOmics Stereo-seq [15]). High-resolution SRT technologies resolve subcellular-level features, either by profiling individual transcript locations (imaging-based SRT) or by capturing multiple transcripts on small spots or nanospheres (sequencing-based SRT). A major challenge of such high-resolution SRT technologies is data sparsity, either due to true biological variations where a gene is not expressed or due to drop-out events caused by technical limitations, which can lead to instability in modelling and downstream analyses [16–21].

Current strategies for overcoming data sparsity include imputing gene expression in the SRT data using a matching reference single-cell transcriptomics (scRNA-seq) dataset or smoothing the gene expression of cells. The data integration strategy is computationally challenging, requiring specific tools to perform imputation [22–28], and is potentially unstable as the reference scRNA-seq data are sparse themselves. In contrast, smoothing is a reference-free approach that generally involves averaging gene expression of neighbouring cells, making it easy to adopt. However, smoothing is insensitive to biological variations, i.e., the complexity of cell type and subtype compositions, often observed in cell neighbourhoods. For example, smoothing over a heterogeneous neighbourhood, where different cell type populations occupy the region, can lead to increased autocorrelation between neighbouring cells, thereby, impacting downstream results [29].

Many existing spatial clustering methods perform local neighbourhood smoothing to utilise the spatial signal in SRT datasets, primarily aiming to partition spots or cells into domains representing anatomical structures in the tissues [30–42]. For this, some methods smooth cell labels using probabilistic approaches [31, 32, 39], whereas others smooth gene expression or low-dimensional embeddings using weighting schemes based on spatial distances [41], histological features [30], or a combination of spatial distances and expression similarity [33]. Generally, these approaches apply a common smoothing parameter to all spots or cells, which determines the balance between self-expression and neighbourhood information; thereby, assuming that all neighbourhoods have a similar level of homogeneity [30–32, 34, 39]. Other methods, like SpaGT, use dynamic spatial graphs and expression similarity to perform smoothing that scales for each cell [43]. However, this approach still relies on pairwise expression similarity and distances, which often struggle when applied to sparse datasets such as high-resolution imaging-based SRT datasets, potentially leading to over-smoothing in highly heterogeneous regions. While these approaches are very effective at domain detection, their applicability to high-resolution SRT datasets to identify subtle cell states is limited. Therefore, methods capable of using the cellular composition of local neighbourhoods in an adaptive manner are required for identification of spatially-informed cell states in high-resolution SRT datasets.

To address this gap, we developed ClustSIGNAL (*<u>Clust</u>*ering of *<u>S</u>*patially-*I*nformed *<u>G</u>*ene expression with *<u>N</u>*eighbourhood-*<u>A</u>*dapted *<u>L</u>*earning), a clustering method that identifies spatially-informed cell states in high-resolution SRT datasets. Our method uses an unsupervised approach to measure each cell’s neighbourhood complexity in terms of entropy, where cells residing in more heterogeneous neighbourhoods have higher entropy compared to cells in more homogeneous neighbourhoods. The entropy values further guide the weights given to each neighbour in a cell’s neighbourhood and the amount of smoothing performed per cell, with low entropy cells being effectively smoothed over a larger neighbourhood and high entropy cells smoothed over a smaller region (see **Methods** section). Here, we used four simulated SRT datasets to assess the robustness of our method parameters and its clustering stability, to compare our adaptive smoothing approach with other smoothing scenarios (no smoothing or complete smoothing of neighbourhood), and to test our method’s applicability to varying levels of technical issues (sparsity, segmentation errors) commonly associated with high-resolution SRT datasets. To demonstrate the performance and scalability of our method compared to some of the existing spatial omics clustering approaches (BANKSY [44], BASS [34], SpatialPCA [45], SpaGCN [30], STAGATE [33], GraphST [36]), we compared the clustering concordance of the methods applied to six real-world datasets generated from various high-resolution SRT technologies (SeqFISH mouse embryo [46], MERFISH mouse hypothalamus preoptic region [47], 10x Xenium breast cancer [48], CosMx SMI lung cancer [9], Stereo-seq ovarian cancer [49], and 10x VisiumHD colon cancer [49]). Finally, we assessed the biological relevance of the clusters obtained from applying ClustSIGNAL to the real-world datasets.

## Results

### ClustSIGNAL uses a neighbourhood complexity-based adaptive smoothing approach for spatially-informed cell-state clustering

We developed ClustSIGNAL, a spatial omics clustering method, for the analysis of high-resolution SRT datasets, which often have very sparse measured expression (**Suppl Fig. S1**). ClustSIGNAL utilises neighbourhood information to stabilise the sparse expression of each cell. For this, it first assesses the neighbourhood complexity (i.e., neighbourhood heterogeneity level) of each cell in an unsupervised manner and then adaptively smooths the cells. This allows ClustSIGNAL to balance between using sufficient information from a homogeneous neighbourhood to stabilise cell expression *versus* refraining from using too much information from a heterogeneous region to avoid over-smoothing (**Suppl Fig. S1**). Briefly, ClustSIGNAL relies on an initial clustering and sub-clustering of the dataset to generate a set of clusters (*initial clusters* and *initial subclusters*) that represent groups of cells with very similar expression patterns (**Fig. 1a**). For each cell, its neighbourhood is then defined by identifying a fixed number of nearest neighbours (default 30 nearest neighbours) in the physical space. The initial subclusters are used to calculate entropy of each cell’s neighbourhood, as an unbiased assessment of neighbourhood composition and heterogeneity. For a given cell, its neighbourhood has low entropy if most of the neighbours belong to the same initial subcluster as the cell, i.e., the cell resides in a more homogeneous region, with zero entropy implying completely homogeneous neighbourhoods. These cell-specific entropy values are used to generate weights for each cells’ neighbourhood, followed by a weighted, adaptive smoothing of gene expression (**Fig. 1b,c**; see **Methods** section). Notably, low entropy, which is associated with more homogeneous neighbourhoods, generates similar weights across all neighbours leading to smoothing over a larger number of the neighbours. In contrast, high entropy, which defines heterogeneous neighbourhoods, generates larger weights for closer neighbours and the weights decay rapidly with increasing distance from the index cell; thereby, leading to smoothing over fewer neighbours. In this way, adaptive smoothing embeds cell-specific spatial information and neighbourhood composition into the expression of each cell. Finally, a clustering method is applied to the adaptively smoothed expression to identify spatially-informed clusters (**Fig. 1c**; see **Methods** section).

**Fig. 1:**
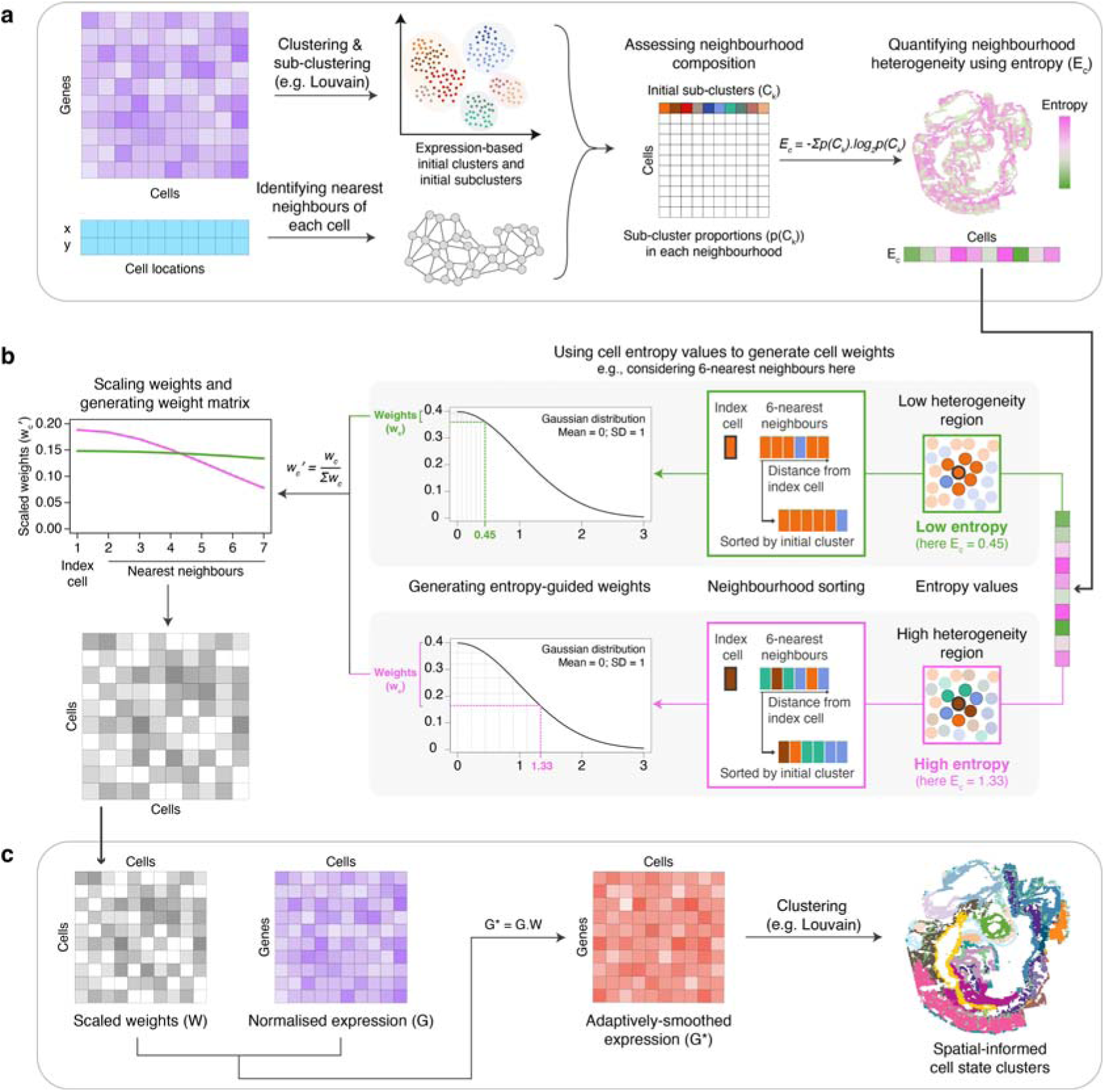
Schematic overview of ClustSIGNAL method. **(a)** *Measuring neighbourhood composition.* A non-spatial clustering approach is used to group the cells into initial clusters and initial subclusters (*C_k_*) based on their gene expression. Following this, each cell’s neighbourhood is defined based on a fixed neighbourhood size (default *N* = 30). The proportion of initial sub-clusters (*p*(*C_k_*)) is assessed in each neighbourhood and used to calculate Shannon’s entropy (*E_c_*) per cell as a measure of its neighbourhood composition. **(b)** *Generating cell-specific entropy-guided weights.* Cell-specific entropy values directly measure neighbourhood heterogeneity. Examples show a low heterogeneity neighbourhood (green outlines and values), where most cells belong to the same initial cluster, and a high heterogeneity neighbourhood (pink outlines and values), where the initial cluster population is more diverse. Prior to weight generation, the neighbouring cells are sorted so that neighbours belonging to the same initial cluster as the index cell are pulled closer to the index cell. Then, for each index cell, its entropy value (*E_c_*) is used to create an entropy sequence vector by uniformly partitioning the range between 0 and *E_c_*. Densities from a Gaussian or exponential distribution (default Gaussian) corresponding to the entropy sequence vector are selected as neighbourhood weight vector (**w***_c_*), which is further scaled (**w** *_c_*) by the sum of its respective weight values. The weights are directly impacted by the entropy value used to generate them, such that low entropy values (low heterogeneity regions) produce similar weights across the neighbourhood (green line in plot), whereas high entropy values (high heterogeneity regions) generate rapidly decaying weights across the neighbourhood (pink line in plot). **(c)** *Adaptively smoothing gene expression and spatial clustering.* Scaled weights are combined into a cell-by-cell weight matrix (**W**), which is used to adaptively smooth the normalised gene expression (*G*) followed by clustering to generate spatially-informed cell state clusters.

ClustSIGNAL has six parameters associated with smoothing or clustering that can impact the final clustering output: neighbourhood size *N*, neighbourhood sorting, kernel type, kernel spread, graph clustering neighbourhood size *k*, community detection method. To select optimal values for these parameters, we rigorously tested various combinations of the parameter values on four simulated datasets of varying spatial structures (Patched, Gradient, Complex, and Uniform; see **Methods** section) with ground truth labels (**Fig. 2a**); parameter combinations were iteratively assessed on each simulated dataset 20 times. We found cell neighbourhood size *N* = 30, Gaussian kernel with *spread* = 0.3 or exponential kernel with *spread* = 5, graph clustering neighbourhood *k* = 10, Louvain community detection [50], and neighbourhood sorting to consistently perform well across all four simulated datasets (**Fig. 2b,c, Suppl Fig. S2**). Although these parameters can take external input from users, we provided optimal values as default or recommended values in the ClustSIGNAL R package, making our flexible and package easy-to-use.

**Fig. 2:**
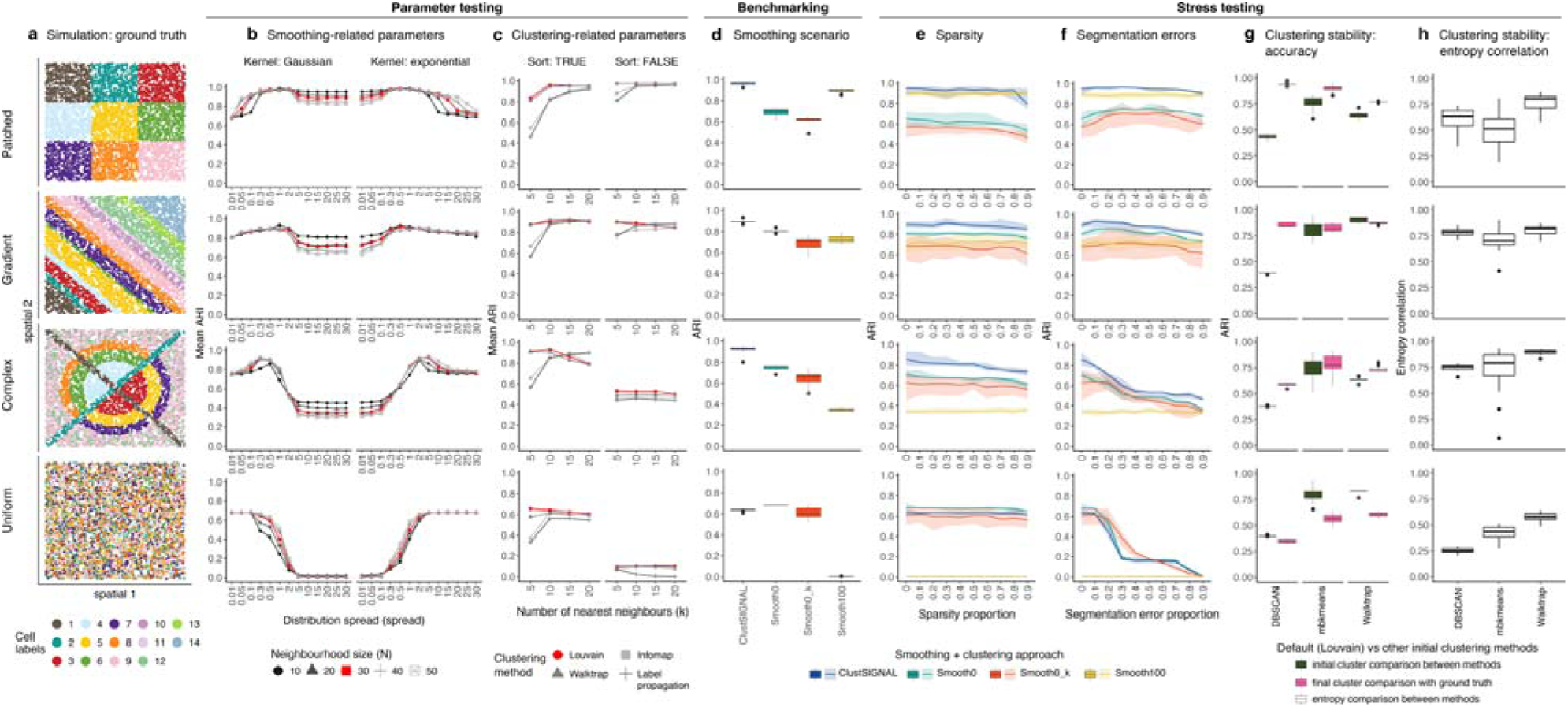
ClustSIGNAL’s adaptive smoothing approach is robust. **(a)** Spatial plots of four simulated datasets showing varying degree of spatial structure, with strong (Patched), intermediate (Gradient and Complex), and no (Uniform) spatial segregation of cell labels (see **Methods** for description of simulated data). **(b, c)** Parameters associated with **(b)** adaptive smoothing, such as neighbourhood size determined by number of nearest neighbours (*N* ; line colours), kernel type (Gaussian, exponential), and kernel spread governed by standard deviation of Gaussian kernel or rate of exponential kernel (x-axis), and **(c)** clustering, such as number of nearest neighbours (*k*) in graph clustering (x-axis) and community detection method (Louvain, Walktrap, Infomap, Label propagation; line colours) were assessed on simulated data to identify optimal values. Other parameters, such as whether or not to sort the neighbourhood, were included with clustering parameters. Optimal values were selected based on highest mean adjusted Rand index (ARI) and number of clusters identified **(Suppl Fig. S2)** in the four simulated datasets and used as default in the R package. **(d)** ClustSIGNAL was benchmarked against: Smooth0, no smoothing; Smooth0_k, no smoothing with the number of clusters set to the number of clusters identified by ClustSIGNAL; and Smooth100, complete smoothing, followed by clustering (**Suppl Fig. S3**). ARI was used to measure clustering performance by comparing cluster labels to the ground truth. **(e, f)** ClustSIGNAL’s capacity to cope with increasing levels (x-axes: 0-90%) of **(e)** sparsity and **(f)** segmentation errors was determined through clustering performance measurement. Comparison was also made between ClustSIGNAL and clustering based on other smoothing approaches (Smooth0, Smooth0_k, Smooth100). **(g, h)** Clustering stability was evaluated by using different initial clustering methods (Louvain, DBSCAN, mbkmeans, Walktrap) in ClustSIGNAL’s algorithm, and measuring **(g)** clustering concordance and **(h)** entropy correlation. **(g)** ARI was measured between (i) initial clusters generated by default (Louvain) *vs* other clustering methods (DBSCAN, mbkmeans, Walktrap) and (ii) final ClustSIGNAL clusters *vs* ground truth. **(h)** Correlation between entropy values generated using initial clusters from default (Louvain) *vs* other clustering methods (DBSCAN, mbkmeans, Walktrap) was also measured.

### ClustSIGNAL’s adaptive smoothing approach is robust

The adaptive smoothing component, together with measured entropy, governs how the spatial information is used by ClustSIGNAL for each cell. To assess the usefulness and limitations of our adaptive smoothing approach, we compared ClustSIGNAL with clustering based on three other smoothing scenarios, i.e., no smoothing (Smooth0), no smoothing with known cluster numbers (Smooth0_k), and complete smoothing (Smooth100), on four simulated datasets with varying levels of spatial structure, decreasing from Patched, Gradient, Complex, to Uniform (**Fig. 2d, Suppl Fig. S3**). Smooth0 and Smooth0_ k, where k represented the number of clusters identified by ClustSIGNAL, were the non-spatial approaches that used only gene expression, whereas Smooth100 was used as another spatial approach that used both gene expression and spatial context. Smooth0_ k was included to assess if a non-spatial approach can partition spatial data in the same way or better than a spatial approach (ClustSIGNAL), given the same number of clusters. In Smooth100, smoothing was performed across all neighbours without any weighting, and so all information was used from the neighbourhood.

#### Resilient to variation in spatial organisation of tissues

We benchmarked ClustSIGNAL against Smooth0-, Smooth0_k-, and Smooth100-based clustering using adjusted Rand index (ARI) to measure clustering concordance. ClustSIGNAL performed consistently better than non-spatial (Smooth0 and Smooth0_k) and spatial (Smooth100) smoothing approaches when applied to simulations with spatial structure, i.e., Patched, Gradient, and Complex (**Fig. 2d**). In Uniform simulation that lacks spatial structure, ClustSIGNAL performance was still comparable to that of the non-spatial clustering approaches (Smooth0, Smooth0_k). Moreover, the loss of spatial structure from Patched to Uniform did not affect ClustSIGNAL accuracy as severely as Smooth100-based spatial clustering, highlighting its adaptability to change in spatial structure due to its selective uptake of neighbourhood information (**Fig. 2d**).

#### Overcomes high levels of data sparsity

We tested the robustness of ClustSIGNAL to increasing levels of data sparsity (0–90% sparsity) in four simulated datasets (Patched, Gradient, Complex, Uniform) and compared its performance against Smooth0-, Smooth0_k-, and Smooth100-based clustering. ClustSIGNAL performance decreased very slowly with increase in sparsity level, and was consistently better than that of other spatial (Smooth100) and non-spatial (Smooth0, Smooth0_k) approaches across most levels of sparsity in Patched, Gradient, and Complex simulations that have some spatial structure (**Fig. 2e**). While Smooth100 was unaffected by sparsity, due to its strategy of averaging over all of the neighbourhood expression, its performance dropped dramatically with decrease in spatial structure, from Patched (high concordance) to Uniform (poor concordance) simulations (**Fig. 2e**). This showed that ClustSIGNAL’s adaptive uptake of neighbourhood information was sufficient to boost gene expression and overcome data sparsity.

#### Robust to increase in segmentation errors

We built ClustSIGNAL specifically for analysis of high-resolution SRT data, where sub-cellular expression is profiled, and generating cell-level counts requires cell segmentation. However, cell segmentation is a challenging problem in SRT, and errors during this process can lead to misallocation of transcripts to incorrect cells [51]. We tested the capacity of ClustSIGNAL to cope with increasing segmentation errors (0–90% segmentation errors) in four simulated datasets (Patched, Gradient, Complex, Uniform) and compared its performance with that of Smooth0-, Smooth0_k-, and Smooth100-based clustering. ClustSIGNAL performance decreased only slightly in Patched and Gradient simulations with increase in segmentation errors, but decreased considerably in Complex and Uniform simulations. However, ClustSIGNAL cluster concordance with ground truth labels was still higher than that of other spatial (Smooth100) and non-spatial (Smooth0, Smooth0_k) approaches in Patched, Gradient, and Complex simulations that have some spatial structure (**Fig. 2f**). In Uniform simulation that lacks any spatial structure, all methods performed poorly with ARI dropping to *<*0.01 at 90% segmentation error. As before, Smooth100 was unaffected by increase in segmentation errors, due to its averaging of information across a large number of nearest neighbours (*N* = 30); however, change in spatial structure greatly reduced its performance. ClustSIGNAL’s adaptive smoothing approach was generally robust to increasing segmentation errors in spatial data, with ARI ranging from 0.5-0.9 at 90% segmentation errors across Patched, Gradient, and Complex simulations.

#### Leads to stable clustering

The first step in ClustSIGNAL algorithm is to generate initial cluster and initial subcluster labels from clustering and sub-clustering of gene expression. The initial subclusters are used to measure each cell’s neighbourhood composition. To assess the stability of ClustSIGNAL clustering, we examined the impact of changes in initial cluster or initial subcluster labels on the clustering outcome by using different initial clustering methods, such as Louvain (default in ClustSIGNAL), DBSCAN [52], mbkmeans [53, 54], or Walktrap [55] on four simulated datasets (Patched, Gradient, Complex, Uniform) (see **Methods** section). This showed low concordance between initial clusters from DBSCAN and Louvain, highlighting the marked differences between the density-based *vs* community detection approach. In contrast, the initial clusters from mbkmeans’ centroid-based approach and Walktrap community detection were comparable to those generated by Louvain method in these simulations, based on their high concordance. Despite the inconsistencies in the initial cluster labels generated by DBSCAN, the performance of the final clusters was comparable across the methods and spatial simulations (Patched, Gradient, Complex) highlighting ClustSIGNAL’s capacity to perform stable clustering irrespective of the choice of initial clustering method (**Fig. 2g**). We also assessed ClustSIGNAL’s approach by comparing the entropy values generated using Louvain initial clustering *vs* those obtained from DBSCAN, mbkmeans, and Walk-trap initial clustering. High correlation between these entropy values in Patched, Gradient, and Complex simulations showed that ClustSIGNAL’s entropy-based approach leads to robust spatial clustering (**Fig. 2h**).

Overall, the simulation analysis showed that the adaptive smoothing-based approach performs better clustering than non-spatial (Smooth0 and Smooth0_k) and other spatial (Smooth100) approaches when applied to spatial data. ClustSIGNAL is robust to changes in spatial structure of samples and can cope with increasing levels of data sparsity and segmentation errors. Moreover, ClustSIGNAL performs stable clustering, with changes in initial clustering method having minor effects on the clustering outcome.

### ClustSIGNAL performs multi-sample spatial clustering in a memory efficient manner

ClustSIGNAL uses an adaptive smoothing approach to generate spatially augmented gene expression and is capable of multi-sample analysis that precludes the need for aligning cluster labels post-clustering. We assessed the performance of ClustSIGNAL on six real-world datasets of varying sizes and from different technologies (**Suppl Table S1**). The datasets included (i) seqFISH mouse embryo (ME) data from Lohoff *et al* [46] containing 351 genes and 52,568 cells in 3 samples, (ii) Xenium breast cancer (BR) data from Janesick *et al* [48] containing 313 genes and 274,501 cells in 2 samples, (iii) CosMx lung cancer (LC) data from He *et al* [9] containing 960 genes and 771,236 cells in 8 samples, (iv) MERFISH mouse hypothalamus (MH) preoptic region dataset from Moffitt *et al* [47] containing 135 genes and 874,756 cells in 181 samples, (v) Stereo-seq human ovarian cancer (OC) data from Ren *et al* [49] containing 31,585 genes and 284,984 cells in one sample, and (vi) VisiumHD human colon adenocarcinoma (COAD) data from Ren *et al* [49] containing 18,042 genes and 520,382 bins in one sample. We compared the performance of ClustSIGNAL with six other spatial clustering methods: BANKSY [44] adaptively uses neighbourhood information to perform spatially-informed cell state or domain clustering; BASS [34] uses a probabilistic approach for multi-sample spatial clustering, including cell type and domain clustering; SpatialPCA [45] performs spatial dimension reduction to generate a joint embedding of gene expression and spatial information that can be used for clustering; SpaGCN [30] uses a graph convolutional network to integrate spatial information and gene expression, followed by clustering; STAGATE [33] applies an attention mechanism to adaptively aggregate neighbourhood information into an embedding used for clustering; and GraphST [36] uses contrastive learning to generate a refined joint embedding for spatial clustering (see **Methods** section). We evaluated method performance by measuring cluster concordance with prior annotations, memory usage, and runtime (**Fig. 3, Suppl Figs. S4-S26, Suppl Table S1**).

**Fig. 3:**
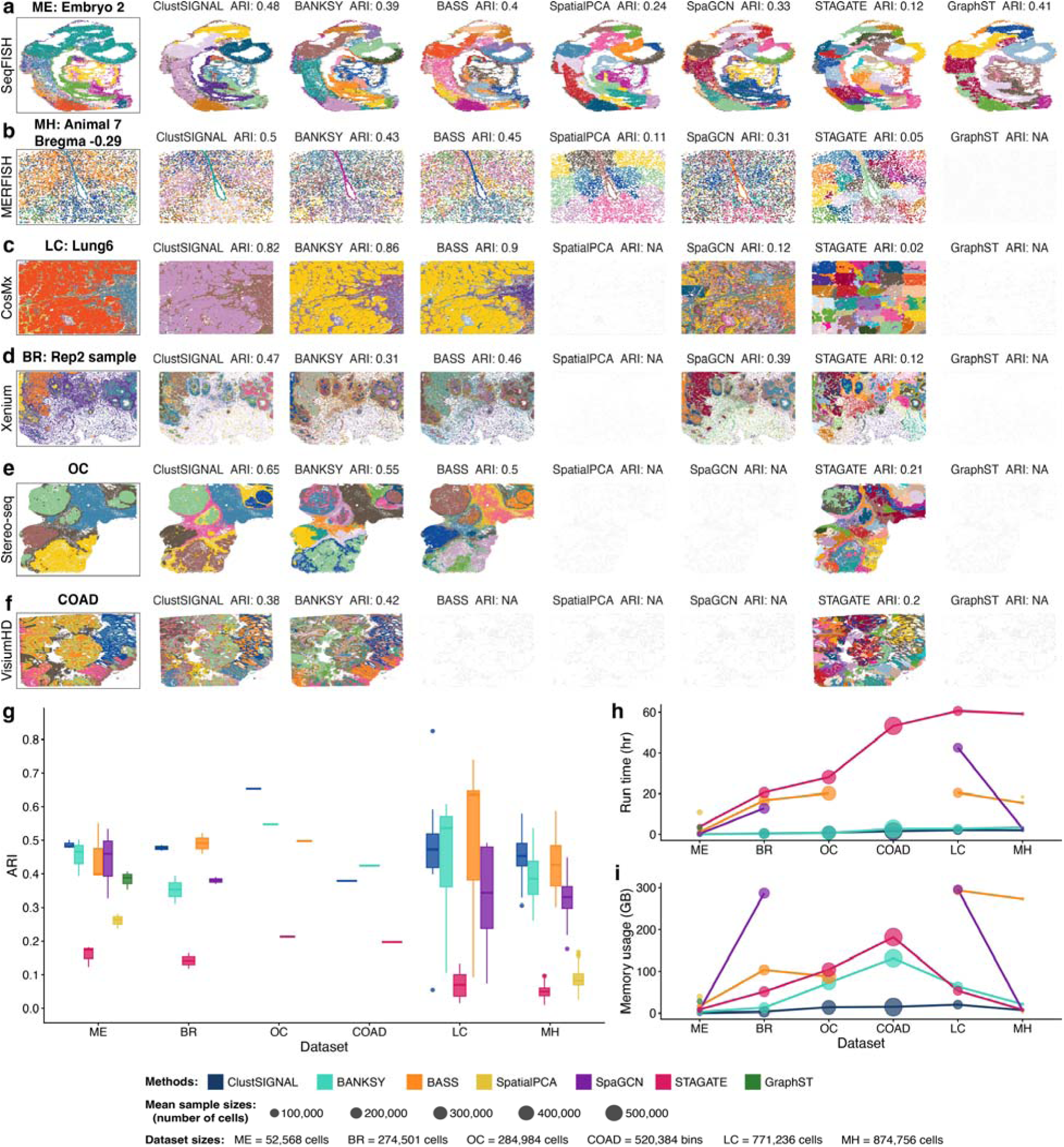
Benchmarking ClustSIGNAL with other spatial clustering methods. **(a-f)** Spatial plots comparing prior annotations (in box; ME: Embryo 2, MH: animal 7 Bregma -0.29, LC: Lung6, BR: Rep2 sample, OC, and COAD) with clusters generated by ClustSIGNAL, BANKSY, BASS, SpatialPCA, SpaGCN, STAGATE, and GraphST performed on **(a)** seqFISH mouse embryo (ME) dataset [46], **(b)** MERFISH mouse hypothalamus (MH) preoptic region dataset [47] **(c)** CosMx lung cancer (LC) dataset [9], **(d)** Xenium breast cancer (BR) dataset [48], **(e)** Stereo-seq ovarian cancer (OC) dataset [49], and **(f)** VisiumHD colon cancer (COAD) dataset [49]. Adjusted rand index (ARI) calculated by comparing cluster labels with prior annotations are shown above each method spatial plot. Spatial plots of all method clusters in the remaining samples from the datasets are also provided **(Suppl Figs. S4-S26)**. **(g)** Overall method performance was measured using the mean of ARI calculated for each sample in a dataset. **(h, i)** Method efficiency was assessed based on **(h)** runtime and **(i)** memory usage on the six datasets organised along the x-axis in an increasing order of dataset size (total cells or bins). All methods were run on the same high-performance computing cluster using only one CPU.

The performance of ClustSIGNAL was consistently better or comparable to that of the other methods on all six datasets (**Fig. 3, Suppl Figs. S4–S26, Suppl Table S1**). ClustSIGNAL had the highest mean ARI in ME, MH, and OC among all methods, whereas in BR, COAD, and LC, its clustering performance (mean ARI) was comparable to that of BASS (BR: 0.48 *vs* 0.49; LC: 0.47 *vs* 0.55) and BANKSY (COAD: 0.38 *vs* 0.42). Notably, ClustSIGNAL used the least amount of memory among all methods on all six datasets, with a maximum of 20 GB memory used to process LC dataset with ∼ 800,000 cells from 8 large samples (*>*70,000 cells per sample) (**Fig. 3i, Suppl Table S1**). ClustSIGNAL’s runtime was also the best on the smallest (ME: ∼ 50,000 cells) as well as the largest (COAD, LC, MH: ∼ 500,000 to ∼ 900,000 cells) datasets, with BANKSY performing faster analysis on BR and OC datasets (∼ 300,000 cells) (**Fig. 3h, Suppl Table S1**). Surprisingly, only ClustSIGNAL, BANKSY, and STAGATE were able to perform clustering on all datasets within the 400 GB memory and 96 hours runtime limits we applied (**Fig. 3, Suppl Table S1**), highlighting their applicability to atlas-level datasets. However, STAGATE’s clustering performance was very poor (mean ARI ≤ 0.2) on all datasets, likely due to comparison between the spatial domains (mixture of cell types) that STAGATE was built to identify *vs* prior annotations that represent distinct cell types. While SpatialPCA was unable to process BR, OC, COAD, and LC datasets containing 300,000–800,000 cells, it was able to perform clustering on MH data with ∼ 900,000 cells (**Fig. 3, Suppl Table S1**). We speculate that this was due to the large number of cells per sample in the datasets rather than the overall dataset sizes. BR, OC, COAD, and LC datasets each contain one or more samples with *>*70,000 cells per sample, compared to MH dataset that has 181 samples containing *<*6000 cells per sample, suggesting that SpatialPCA requires substantial computational resources to handle datasets with very large samples.

Overall, ClustSIGNAL is highly scalable to atlas-level datasets with large samples, generating clusters with high concordance to prior annotations (**Fig. 3, Suppl Figs. S4–S26, Suppl Table S1**). Additionally, ClustSIGNAL can perform multi-sample analysis without relying on spatial alignment of the samples, as it searches for neighbours in individual samples and combines the entropy values from all samples for the following steps (**Suppl Fig. S27, S28**). For samples with strong batch effects, ClustSIGNAL supports integration of the samples by employing Harmony integration internally, similar to BASS (see **Supplementary Material**).

### ClustSIGNAL performs spatially-informed cell-state clustering by adaptively incorporating spatial information

We examined the clusters obtained from applying ClustSIGNAL to ME and MH datasets to assess their biological relevance, specifically clusters that represented subgroups of prior annotations. In ME data, ClustSIGNAL identified 22 distinct clusters across 3 mouse embryo samples (Embryo1, 2, 3), with some clusters representing subgroups of prior annotations, such as forebrain/midbrain/hindbrain (clusters 4, 10, 12, 13), gut tube (clusters 17, 22), spinal cord (clusters 9, 19), and neural crest (clusters 14, 21) (**Fig. 4a,b, Suppl Fig. S29**). We investigated the gene expression and spatial patterns of the forebrain/midbrain/hindbrain clusters (4, 10, 12, 13) and found that they expressed distinct marker genes (**Fig. 4c**). Of the four clusters, only cluster 10 cells expressed *Hoxb1* gene known to be involved in the development of hindbrain [56], clusters 12 and 13 had genes associated with regulation and development of forebrain (*Dlk1* [57], *Fezf1* [58], *Six3* [59–62], and *Lhx2* [63]), and cluster 4 had high expression of *En1* gene involved in the survival and maturation of midbrain dopaminergic neurons [64, 65] (**Fig. 4c**). A pathway enrichment analysis of the top marker genes of these clusters supported our inference that clusters 12 and 13 contained genes associated with forebrain development and regionalisation, whereas cluster 4 contained marker genes related to midbrain development (**Fig. 4d**). While we did not observe any hindbrain-associated pathways enriched in cluster 10, a few marker genes were associated with midbrain development suggesting an expression continuum across the midbrain and hindbrain regions given their spatial adjacency. The association of these clusters with specific brain regions was also evident from the high expression of their marker genes in specific parts of the brain region on the tissues (**Fig. 4e**). While both clusters 12 and 13 contained forebrain cells, we observed cluster 12 only in Embryo1 and Embryo3 and cluster 13 only in Embryo2, with cluster 13 showing a much higher expression of *Setd2* gene associated with the development of forebrain regions [66] (**Fig. 4a,c**). ME data contained the X-chromosome gene *Xist*, which had no expression in Embryo2 tissue that was extracted from a male mouse. Re-running ClustSIGNAL after removing the *Xist* gene from ME data did not affect the partitioning of forebrain cells, suggesting that the cluster separation was likely not driven by biological sex. Similarly, clusters representing subgroups of gut tube (clusters 17, 22), spinal cord (clusters 9, 19), and neural crest (clusters 14, 21) had distinct markers and occupied specific spatial regions indicating that they represented different cell subtypes (**Suppl Fig. S29**). A UMAP projection with pseudotime trajectory overlay also showed that ClustSIGNAL captures cell states, which represent continuous gene expression patterns, without blur-ring the boundary between the distinct cell types (**Fig. 4f**). This demonstrates that ClustSIGNAL’s entropy-guided adaptive smoothing preserves underlying biological structure and avoids over-smoothing.

**Fig. 4:**
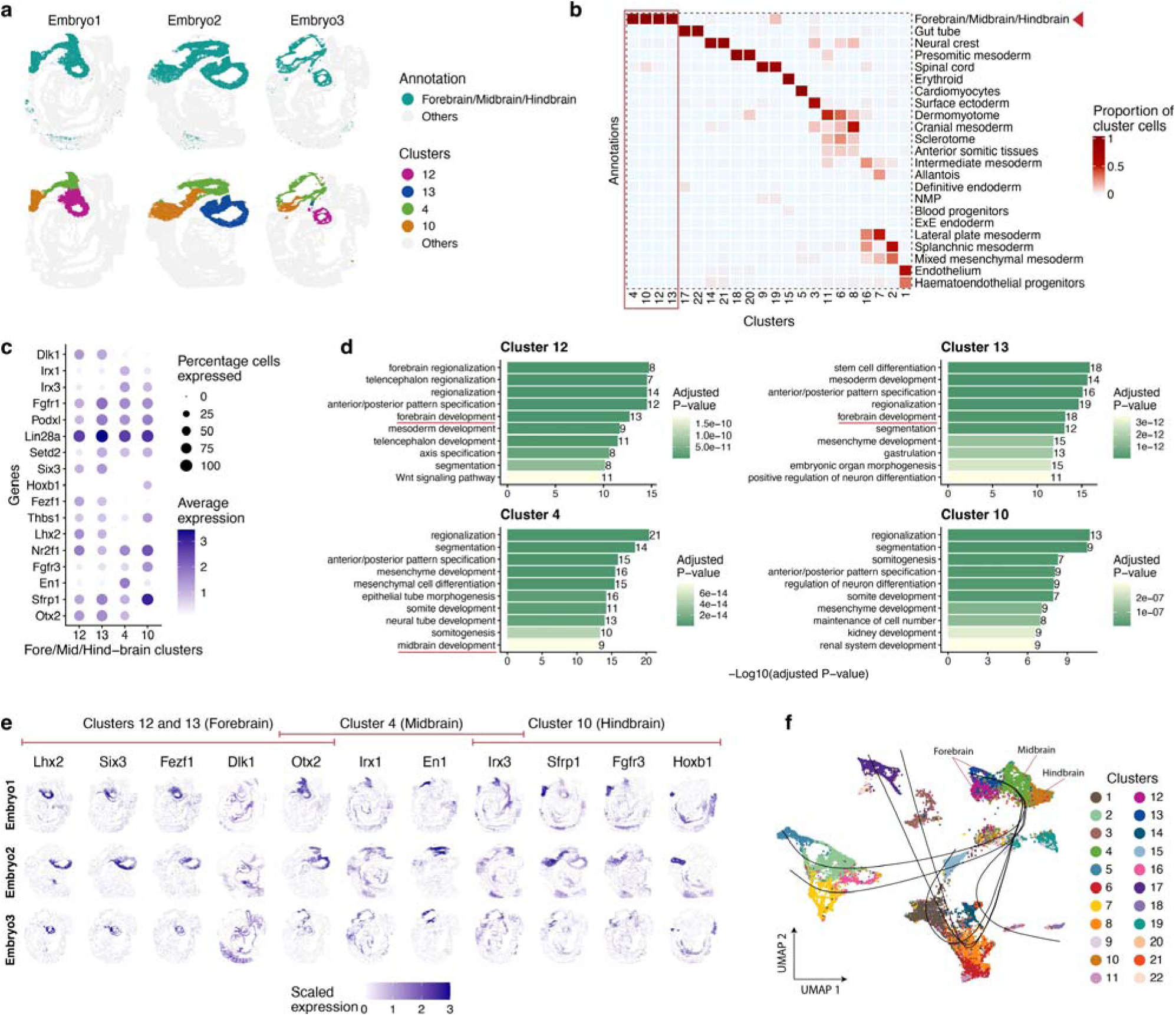
ClustSIGNAL identifies brain region subtypes in mouse embryo development (ME) dataset. **(a)** Spatial plots of the three ME samples (Embryo1, Embryo2, Embryo3) from Lohoff *et al* [46] showing forebrain/midbrain/hindbrain annotation (top row) and ClustSIGNAL clusters associated with the brain region (bot-tom row). **(b)** Heatmap showing proportion of ClustSIGNAL cluster cells (columns) matching the prior annotations (rows) in ME data. **(c)** Dot plot of top 10 marker genes from each of the brain-associated clusters (4, 10, 12, 13). **(d)** Bar plots showing top 10 biological pathways (BP ontology) significantly enriched in the brain clusters based on differentially-expressed (DE) genes (*log*_2_*FC >* 0.2*, FDR <* 0.05) in each cluster. Brain development associated pathways are highlighted in the plots (red underline), except for cluster 10. Only pathways associated with at least 5 DE genes were considered. **(e)** Spatial plots showing scaled normalised expression of the marker genes **(c)** associated with brain region clusters from ME data. **(f)** UMAP coloured by ClustSIGNAL clusters, with an overlay of inferred developmental lineages. The branching pseudotime trajectories highlight the continuous biological gradient that correctly maps the anatomical structure of the brain region (Forebrain, Midbrain, Hindbrain).

In the 181 samples from MH data, ClustSIGNAL generated 12 clusters, some of which represented subgroups of Inhibitory (clusters 5, 6, 8) and Excitatory (clusters 3, 10) neurons (**Fig. 5a,b**). On investigating the gene expression and spatial pattern of Inhibitory and Excitatory clusters, we observed that they expressed distinct marker genes and were spatially organised in similar regions across all tissues from specific bregma slices (**Fig. 5a,c, Suppl Figs. S30, S31**). We further examined the neighbourhood composition of the inhibitory clusters by assessing co-localisation of clusters and found stark differences in their neighbourhood organisation in different tissue slices (**Fig. 5a,d**). For example, in Animal 8 Bregma –0.24 and Animal 7 Bregma –0.14 tissues (i.e., tissue slices from posterior of the bregma), Inhibitory cluster 8 cells mainly co-localised with themselves, likely forming homogeneous micro-domains in the tissue, whereas Inhibitory cluster 5 cells mainly co-localised with other cluster cells, suggesting a more dispersed presence in these tissues (**Fig. 5d**). This was also supported by their spatial organisation in the tissues, where slices from the posterior of bregma (negative bregma slices) were dominated by homogeneous streaks of Inhibitory cluster 8 cells and slices from the anterior of bregma (positive bregma slices) had a diffused dominant population of Inhibitory cluster 5 cells (**Fig. 5a, Suppl Fig. S30**). Similarly, Excitatory clusters showed a distinct co-localisation pattern, where cluster 10 cells mainly co-localised with themselves in all tissues and cluster 3 cells co-localised with a number of other clusters (**Fig. 5a,d, Suppl Fig. S31**). Cell neighbourhood analysis also showed that these Inhibitory and Excitatory clusters were associated with specific cell types (**Fig. 5e**). For example, OD Immature cells were often significantly more associated with Inhibitory cluster 8 than with other clusters, whereas OD mature cells were significantly co-localised with Inhibitory cluster 5 compared to other clusters. Excitatory cells showed distinct association patterns in tissues from posterior (Animal 8 Bregma –0.24, Animal 7 Bregma –0.14) *vs* anterior (Animal 12 Bregma 0.21, Animal 17 Bregma 0.26) of the bregma (**Fig. 5e**). In the posterior tissues, Excitatory cells were associated more with Inhibitory cluster 6 cells than with other clusters, whereas in the anterior tissues, Excitatory cells co-localised significantly more with both Inhibitory (5, 6) and Excitatory (3, 10) clusters compared to all other clusters. We could not assess the impact of these associations through a cell-cell interaction analysis due to the limited number of genes (only 135 genes) in the MH dataset.

**Fig. 5:**
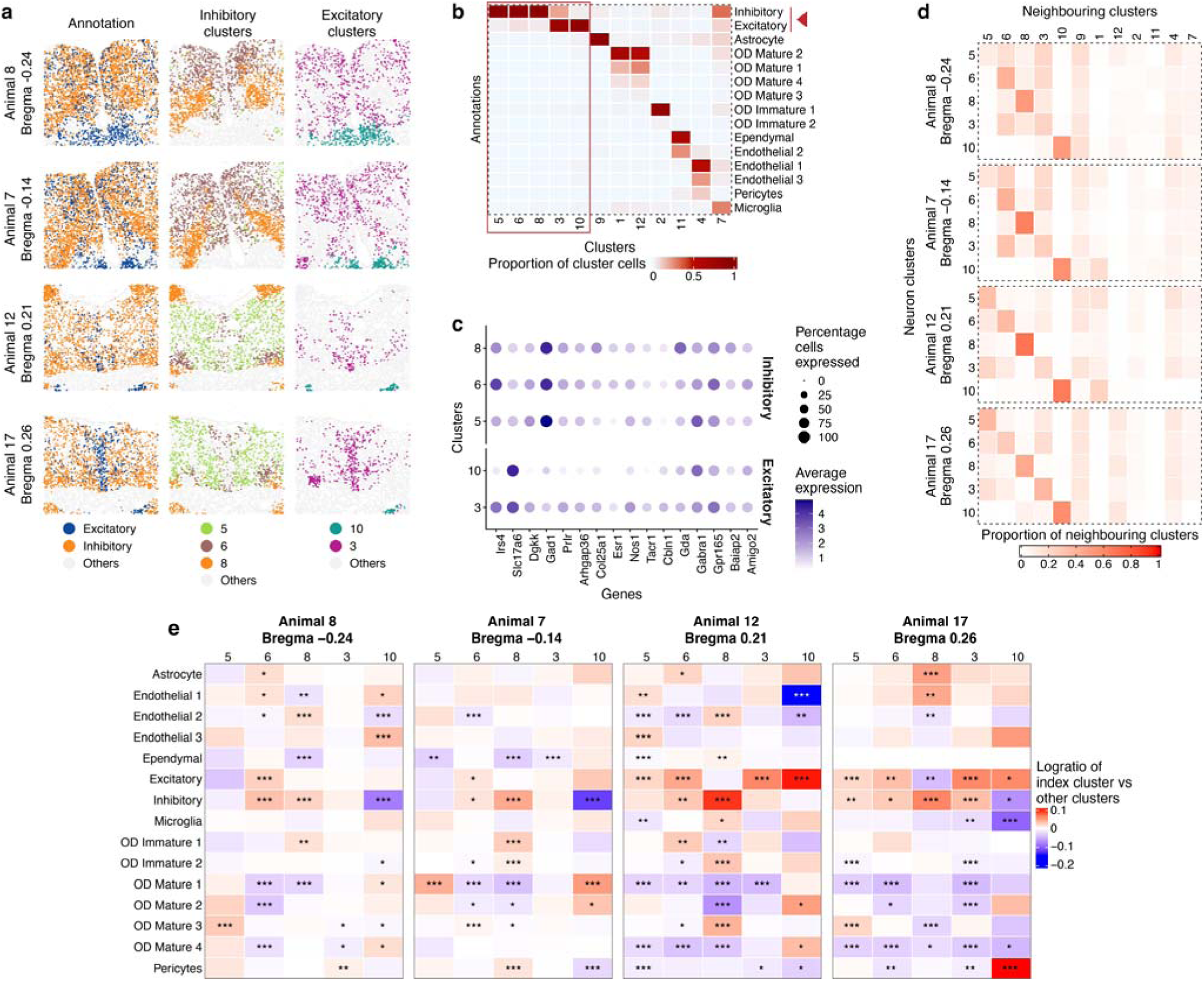
ClustSIGNAL adaptively uses neighbourhood composition to identify neuron subtypes in mouse hypothalamus (MH) dataset. **(a)** Spatial plots of four MH samples (Animal 8 Bregma - 0.24, Animal 7 Bregma -0.14, Animal 12 Bregma 0.21, Animal 17 Bregma 0.26) from Moffitt *et al* [47] showing Inhibitory and Excitatory neuron annotations (left column), clusters associated with Inhibitory neurons (middle column), and clusters associated with Excitatory neurons (right column). Spatial plots showing organisation of Excitatory and Inhibitory clusters in all samples are also provided **(Suppl Fig. S30, S31)**. **(b)** Heatmap showing proportion of ClustSIGNAL cluster cells (columns) matching the prior annotations (rows) in MH data. **(c)** Dot plot of top 10 marker genes from Inhibitory (5, 6, 8) and Excitatory (3, 10) neuron associated clusters. **(d)** Co-localisation heatmaps showing proportion of neuron associated cluster cells (rows) that co-localised with the neighbouring cluster cells (columns) in each sample. **(e)** Heatmap of cell type neighbourhood analysis of Inhibitory (5, 6, 8) and Excitatory (3, 10) neuron clusters in each sample, highlighting their distinct associations with specific cell types. Colour and values indicate whether a neighbouring cell type (rows) is more associated with the index cluster in each column (red, positive values) or other clusters in the dataset (blue, negative values). BH-corrected *P* -values: * *<*0.05; ** *<*0.01; *** *<*0.001.

A similar refinement of annotations was observed when ClustSIGNAL was applied to BR and LC datasets, where some annotations were sub-grouped into clusters with specific marker genes (**Suppl Figs. S32, S33**). These data suggested that ClustSIGNAL identifies cell types and cell subtypes of biological relevance using gene expression and adaptively incorporated neighbourhood information.

## Discussion

In this paper, we introduce ClustSIGNAL, a spatially-informed cell-state clustering method. A key component of ClustSIGNAL is its novel adaptive smoothing approach that relies on each cell’s neighbourhood complexity gathered in an unsupervised manner using neighbourhood entropy, rather than on pairwise expression similarities of neighbouring cells. Our approach leads to selective uptake of local gene expression for each cell depending on its neighbourhood organisation (**Fig. 1**). Therefore, cells in homogeneous neighbourhoods are smoothed over a larger number of neighbours, which stabilises their gene expression, whereas cells in more heterogeneous tissue spaces are smoothed over a smaller number of closest neighbours, which ensures that their distinct gene expression is preserved. In this way, ClustSIGNAL uses smoothing to handle data sparsity without over-smoothing cells that lie in highly heterogeneous regions. It preserves the biological structure of the tissues while generating clusters that represent cell types and cell subtypes.

ClustSIGNAL’s adaptive smoothing approach is robust to technical issues, such as sparsity and segmentation errors, commonly observed during high-resolution SRT data analysis, and leads to stable clustering (**Fig. 2**). We applied ClustSIGNAL to four simulated spatial datasets with varying degrees of spatial structure. ClustSIGNAL’s performance was resilient to increasing levels of sparsity and segmentation errors in the simulations containing spatial structure. A comparison between different types of clustering methods (density-based, centroid-based, community detection) used for initial clustering and sub-clustering of the simulated datasets showed that ClustSIGNAL’s adaptive smoothing approach mitigated the differences in initial clusters and subclusters obtained from these methods and led to stable clustering in simulations with distinct spatial structure.

ClustSIGNAL is highly scalable and can be applied to datasets with multiple samples, with inherent functionality to call Harmony for integrating samples with technical artifacts, such as strong batch effects. It extracts cell-specific neighbourhood information from individual samples and combines the resulting entropy values into a singular framework; therefore, ClustSIGNAL does not rely on spatial alignment of samples for multi-sample analysis (**Suppl Figs. S27, S28**). We built ClustSIGNAL on an existing data structure (SpatialExperiment) and rigorously tested the default values for the parameters provided within the R package. Its application to six real-world datasets from different SRT technologies showed that ClustSIGNAL was scalable to atlas-level datasets, performing fast and memory efficient clustering compared to existing spatial clustering methods (**Fig. 3, Suppl Table S1**).

ClustSIGNAL is applicable to 2D and 3D spatial data from high-resolution spatial omics platforms, including imaging-based and sequencing-based technologies. Its reliance on *n*-dimensional matrix of spatial coordinates for detecting neighbours means that ClustSIGNAL can also be applied to 3D spatial data. While ClustSIGNAL can technically be applied to low-resolution SRT datasets, such as 10x Visium, we do not recommend it; these datasets capture multiple cells per spot, and the resulting clusters would be biologically difficult to interpret. Finally, we developed ClustSIGNAL for clustering high-resolution spatial transcriptomics data, but it can be applied to other high-resolution spatial omics datasets with cell-level information, e.g., spatial proteomics datasets, albeit with compensations for differences in the input data type, such gene counts *vs* intensities (**Suppl Fig. S34**; see **Supplementary Material**).

### Conclusions

ClustSIGNAL provides a novel neighbourhood complexity-based approach for adaptive smoothing of neighbourhoods for tackling data sparsity and performing unsupervised spatially-informed cell-state clustering of high-resolution spatial omics data. Our results show that adaptive smoothing effectively stabilises gene expression in low heterogeneity regions and avoids over-smoothing regions with high heterogeneity, leading to stable clustering that is resilient to sparsity and segmentation errors. As the spatial omics field is moving toward large-scale datasets, there is a necessity for robust and scalable methods. ClustSIGNAL addresses this need through memory-efficient, multi-sample analysis of atlas-level datasets to generate reliable and biologically interpretable clusters from high-resolution omics data.

## Methods

### Implementation of ClustSIGNAL framework

We developed ClustSIGNAL to perform spatial clustering of high-resolution SRT datasets, using a modular approach with five sequential steps: (i) clustering and sub-clustering of normalised gene expression to generate initial clusters and subclusters; (ii) identifying each cell’s *N* nearest neighbours; (iii) calculating entropy of each cell’s neighbourhood from proportions of initial subclusters; (iv) using neighbourhood entropy values to generate cell-specific weight vectors for adaptively smoothing gene expression; and (v) clustering adaptively smoothed gene expression (**Fig. 1, Suppl Fig. S27**). ClustSIGNAL can be run on multiple cores, and is available as an R package on Bioconductor bioconductor.org/packages/clustSIGNAL.

#### Initial clustering and sub-clustering to identify cells with similar transcriptomic profiles

We designed ClustSIGNAL to accept a SpatialExperiment [67] object containing normalised gene expression and spatial coordinates as input. We perform principal component analysis (PCA) using *runPCA()* function from the scater R package [68] to generate a low dimensional embedding. Using this embedding, we cluster the cells with the *TwoStepParam()* function from the bluster R package, which performs a k-means clustering followed by a nearest-neighbour graph community detection (Louvain by default) to generate initial clusters. This two-step clustering approach allows us to improve the computational efficiency of the clustering step by performing a k-means micro-clustering with centres equal to 3000 or 1/5th of the total cells in the dataset, whichever is lower. Each initial cluster is further sub-clustered into initial subclusters using the *TwoStepParam()* function with centres equal to 1 or half of the total cells in the cluster, whichever is higher, for the k-means clustering component. Here, we assume that the initial subclusters represent groups of cells with very similar expression profiles. For multi-sample analysis, we also provide specific parameters (*batch* and *batch_by*) within the ClustSIGNAL package to perform a Harmony [69] integration of gene expression in samples with strong batch effects, prior to initial and final clustering.

#### Detecting and sorting neighbours

We use the *findKNN()* function from the BiocNeighbors R package to identify *N* nearest-neighbours (default *N* = 30) of each cell in the physical space. We refer to the spatial neighbourhood of each cell as region **r***_c_*, with one region per cell in the dataset. We define *i* as a set of cells *c_i_* in region **r***_c_*, i.e. *i* = *{*1, 2*, . . ., N* + 1*}*, where *c*_1_ is the index cell and *c_i_* for *i >* 1 are its neighbours. Within each region **r***_c_*, the neighbouring cells are ordered based on their increasing physical Euclidean distance from the index cell *c*_1_, with the closest neighbours placed adjacent to the index cell in the **r***_c_* vector. By default, we perform an additional step of sorting each region **r***_c_* such that cells belonging to the same initial cluster as the index cell are placed closer to it in the **r***_c_* vector; the order of all other cells in the region remains unchanged. This sorting step allows us to give higher weights to the cells with expression profiles similar to that of the index cell, to ensure that information is shared mainly between similar cells.

#### Measuring neighbourhood heterogeneity as entropy

We measure the composition of each region **r***_c_* by estimating its heterogeneity level using Shannon’s entropy *E_c_*. For this, we use the proportion of initial sub-clusters in each region **r***_c_* to calculate its entropy *E_c_*.

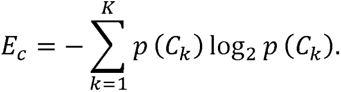

Here, we define *k* as a set of initial subclusters *C_k_*, i.e. *k* = *{*1, 2*, . . ., K}*, where *K* is the total number of initial subclusters identified in the dataset. The proportion of an initial subcluster in a region **r***_c_* is represented by *p*(*C_k_*). The entropy values generated here are cell-specific, ranging from 0 to log of *K* initial subclusters in the dataset, and represent an unsupervised measure of a cell’s neighbourhood complexity. This means that cells with high entropy reside in more heterogeneous regions than cells with lower entropy; cells with *E_c_* = 0 lie in completely homogeneous neighbourhoods. Note that cell entropies *E_c_* are strongly dependent on the neighbourhood size *N* across which neighbourhood composition was measured.

#### Generating entropy-guided weights

In ClustSIGNAL, we use the cell-specific entropy values to determine the “effective” smoothing radius of each cell, and generate cell-specific weight vectors for the neighbourhoods. For this, we first split the range between 0 and the cell’s entropy *E_c_* into equal intervals to generate a uniformly spaced entropy vector **x***_i_* of *N* + 1 length. Biologically, this entropy vector models the decaying spatial gradient of the influence of the neighbouring cells in **r***_c_* of cell *c*.

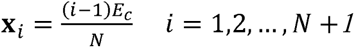

Following this, we apply *dnorm()* or *dexp()* function to the entropy vector **x***_i_* and get a weight vector of Gaussian or exponential distribution densities, respectively (**Fig. 1b**). For each cell, we generate a weight vector **w***_c_* of length equal to the number of cells in region **r***_c_* based on entropy *E_c_* of the index cell. While the Gaussian and exponential distributions are highly related, the weights generated from an exponential kernel show a faster initial decay close to the origin compared to the flatter and smoother weights generated from a Gaussian kernel [70, 71]. This allows the exponential kernel to generate weights with a much stronger local focus, preserving the expression of the index cell.Therefore, we recommend using an exponential kernel (spread ≥5) for datasets from highly heterogeneous tissues.

When using Gaussian distribution,

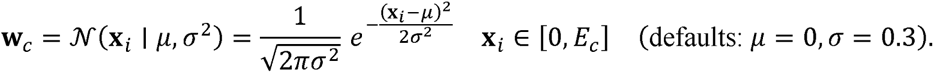

When using exponential distribution,

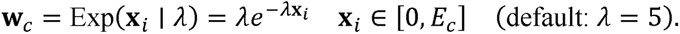

Finally, we scale the weight vector **w***_c_* by the sum of its weight values:

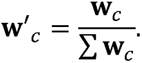

Inherently, the densities in the scaled weight vector **w** *_c_* are arranged in a decreasing order, and are very similar when the input entropy value is very low (**Fig. 1b**). For example, for *E_c_* = 0, all densities in the weight vector have the same value. For high entropy, only the first few densities are very high, and the remaining weight vector can have negligible values (**Fig. 1b**). In ClustSIGNAL, we use this relationship between the entropy values and density weights to adaptively incorporate neighbourhood information into the gene expression during smoothing. Heterogeneous regions have high entropy, which means that ClustSIGNAL gives more weight to cells closer to the index cell and negligible weight to cells farther away; thereby, smoothing over a smaller number of cells. In contrast, homogeneous regions have low entropy, which means that ClustSIGNAL gives similar weights to most cells in these regions, and smooths over a larger number of cells.

#### Adaptively smoothing gene expression

We use the composition of each cell’s neighbourhood to create a modified gene expression matrix with spatial information and neighbourhood complexity incorporated in a cell-specific, adaptive manner. We perform weighted averaging of the normalized gene expression **G** using the weight matrix **W** composed of cell-specific scaled weights **w** *_c_* to obtain adaptively smoothed expression:

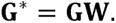

For the adaptively-smoothed expression of each gene *g* in the index cell *c*,

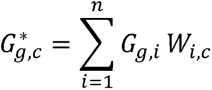

where *G*_g,c_* is the adaptively-smoothed expression of gene *g* in index cell *c*, *n* is the total number of cells in the dataset, *G_g,i_* is the normalised expression of gene *g* in the *i*th cell, and *W_i,c_* is the scaled weight allocated to the *i*th cell in the weight matrix. **W** is a cell-by-cell (*n* x *n*) weight matrix, where the columns are the index cells, the rows are all cells including the neighbouring cells, and values are scaled weights of all cells; cells that are not present in the neighbourhood **r***_c_* of index cell *c* are given zero weights.

#### Clustering adaptively smoothed gene expression

Finally, we cluster the adaptively-smoothed gene expression using the parameters for initial clustering. We first perform PCA to generate low dimensional embedding of the adaptively smoothed data. If batch correction is opted for (*batch* = *TRUE*), we generate a batch-corrected low dimension embedding using Harmony. As before, we cluster the cells using the *TwoStepParam()* function from the bluster R package. This performs a k-means clustering, where we use centres equal to 3000 or 1/5th of the total cells in the dataset, whichever is lower, followed by a graph community detection (Louvain by default) to generate final spatial clusters.

### Datasets: overview and processing

We used six real-world datasets to demonstrate the applicability of ClustSIGNAL to high-resolution SRT data (see **Data availability**). We selected these datasets based on (i) tissue type (developmental, healthy, and disease tissues), (ii) number of samples in the dataset (ranging from 1 to 181), (iii) number of cells in the dataset (ranging from 50,000 to nearly a million cells or bins), and (iv) different SRT technologies (seqFISH, MERFISH, Xenium, CosMx, Stereo-seq, or VisiumHD).

#### Mouse embryo (ME) dataset

Raw and normalized gene expression counts and cell metadata were downloaded from Lohoff *et al* [46], generated using seqFISH technology. SRT data were profiled from 3 sagittal sections of a WT C57BL/6J mouse embryo in E8.5–8.75 stage of embryonic development, with 351 genes and 57,536 cells. Cells with prior annotations as “Low quality” were excluded, and 52,568 cells were used from all analyses.

#### Human breast cancer (BR) dataset

Transcript-level expression data were downloaded from the 10X Genomics website, containing information from 2 tissue section replicates from a formalin-fixed, paraffin-embedded (FFPE) breast cancer (TNM stage T2N1M0) tissue block processed using 10X Genomics Xenium platform. Raw transcript-level data, i.e., transcript locations and cell and nucleus boundaries, were read using MoleculeExperiment R package [72], and cell-level gene counts were obtained by running *countMolecules()* function on the *me* object. Processed data from Janesick *et al* [48] were used to select the 313 panel genes, and exclude the negative control genes, as well as to obtain cell type an-notations for a total of 274,501 cells. Log-transformed normalised gene counts were generated using the *logNormCounts()*function in scater R package.

#### Human lung cancer (LC) dataset

Gene expression information were downloaded from the Nanostring website in the form of a pre-processed Giotto R object from He *et al* [9]. The data were profiled from 5 FFPE tissue of human non-small cell lung carcinoma. Raw counts of 960 genes and metadata of 771,236 cells, extracted from the Giotto object, were stored in a SpatialExperiment R object and used for all analyses. The dataset showed strong batch effects between samples from different patients. Therefore, for data analysis with ClustSIGNAL, batch correction was performed using Harmony internally (*batch* = *TRUE*) based on patient group (*batch_by* = “patient”). For benchmarking, batch correction was not performed for fair comparison with methods that do not perform multi-sample analysis.

#### Mouse hypothalamus preoptic region (MH) dataset

Gene expression counts and cell metadata were downloaded from Moffitt *et al* [47] generated on the MERFISH platform. For this dataset, a total of 181 mouse hypothalamic preoptic tissue slices from 6 social behaviour types in adult C57Bl6/J mice, 16 female and 20 male mice, were profiled with a panel of 155 genes measured in 1,027,080 cells. For consistency, 20 genes profiled using a different technology (non-combinatorial seqFISH technology) and cells with prior annotations as “Ambiguous” were filtered out. A total of 135 genes and 874,756 cells were used for all analyses. Normalised gene counts were generated using the *logNormCounts()* function in scater R package.

#### Human ovarian cancer (OC) dataset

Gene expression counts and cell metadata were downloaded from the SPATCH website as an AnnData object of the Stereo-seq sample from Ren *et al* [49]. The data were profiled from a fresh frozen human ovarian cancer tissue. Raw counts of 32,008 genes in 319,366 cells were extracted from the AnnData object and stored in a SpatialExperiment object. Cells with no annotation or those not profiled with CODEX were removed, and genes with no expression in filtered cells were removed. Total 31,585 genes and 284,984 cells were used for analysis. Normalised gene counts were generated using the *logNormCounts()* function in scater R package.

#### Human colon adenocarcinoma (COAD) dataset

Gene expression counts and 8-*µ*m bin metadata were downloaded from the SPATCH website in the form of an AnnData object of the VisiumHD sample from Ren *et al* [49]. The data were profiled from a FFPE human colon adenocarcinoma tissue. Raw counts of 18,085 genes in 650,432 bins were extracted from the AnnData object and stored in a SpatialExperiment object. Bins with no annotation or those not profiled with CODEX were removed, and genes with no expression in filtered bins were removed. Total 18,042 genes and 520,382 bins were analysed. Normalised gene counts were generated using the *logNormCounts()* function in scater R package.

#### Simulated data analysis

We performed simulated data analysis to assess: (i) proof of concept, (ii) perform parameter tuning to identify optimal values, and (iii) perform stress testing to understand the limits of the method and the contributions of specific method components. To generate a simulated spatial data where the ground truth is known, we needed gene expression that captures the complexity of real data. Therefore, we used gene expression from the single cell transcriptomic (scRNA-seq) atlas of human tissues “Tabula Sapiens” from the CellxGene database (see **Data availability**). We selected cells captured using 10x 3’ v3 assay from specific donors to minimize batch effects — plasma cells and B cells from lymph node of donor TSP14; mesenchymal stem cells and skeletal muscle satellite stem cells from muscle of donor TSP14; hepatocyte from liver of donor TSP14; conjunctival epithelial cells, corneal epithelial cells, and retinal blood vessel endothelial cells from eye of donor TSP21; and kidney epithelial cells from kidney of donor TSP2 were used. Reference gene expression was generated by combining the scRNA-seq data from the above mentioned nine cell types into a SingleCellExperiment object [73] — B cells (4,203), plasma cells (1,127), mesenchymal stem cells (4,842), skeletal muscle satellite stem cells (1,632), hepatocytes (5,393), conjunctival epithelial cells (4,214), corneal epithelial cells (2,657), retinal blood vessel endothelial cells (1,391), and kidney epithelial cells (9,126). Following this, the top 3000 highly variable genes were selected using the scran R package [74].

Next, we generated spatial locations in four patterns with varying degrees of spatial structure — *Patched* had the simplest pattern where each ground truth label, hereafter referred to as “cell label”, was allocated a distinct region and each region had similar number of cells; *Gradient* also had cell labels allotted to distinct regions but there was some disparity between the sizes of the regions and the number of cells allotted to them, *Complex* had distinct regions with specific cell label allotments and regions with no structure where a mixture of cell labels could be found, and *Uniform* represented sample with no spatial structure where all cells from all cell labels were randomly distributed across the whole sample space (**Fig. 2a**). Gene expression was assigned to all cells belonging to a cell label in two ways: (i) randomly selecting cells from one of the scRNA-seq reference cell types and using their gene expression, or (ii) randomly selecting cells from two neighbouring reference cell types and using their averaged gene expression. With the latter strategy, we incorporated spatial information into the data based on the principle of spatial autocorrelation, which assumes that neighbouring cells likely share some expression similarity, and the concept of a biological expression continuum. For example, if cells belonging to cell label X exist between cells belonging to cell labels A and B, we randomly select two cells labelled A and B, average their gene expression, and assign it to a random cell labelled X. A table showing the combination of reference cell types used for simulated cell labels in each pattern is provided (**Suppl Table S2**).

### Parameter tuning

We tested a total of six parameters from ClustSIGNAL to identify optimal values to use as default. First, we assessed three parameters associated with incorporation of spatial information, namely number of nearest neighbours *N* (10, 20, 30, 40, 50), type of distribution kernel (Gaussian, exponential), and kernel spread (0.01, 0.05, 0.1, 0.3, 0.5, 1, 2, 5, 10, 15, 20, 25, 30) on the four simulated datasets (Patched, Gradient, Complex, Uniform), for a total of 520 combinations. Each combination was assessed iteratively 20 times for robustness. We selected optimal values for *N*, kernel, and spread and used them with the remaining three parameters for a second batch of parameter testing. For this, the two parameters associated with clustering, i.e., number of k-means centres (5, 10, 15, 20) and community detection method (Louvain, Walktrap [55], infomap [75], label propagation [76]), and parameter for whether to perform neighbourhood sorting prior to adaptive smoothing (True, False) were assessed on the four simulated data (Patched, Gradient, Complex, Uniform), for a total of 128 combinations — 20 iterations of each combination were run for robustness. The optimal values were selected based on high clustering concordance (mean ARI) measured using the *ARI()* function from the aricode R package, and the least amount of variation in the average number of clusters generated compared to ground truth in all four simulated datasets (**Fig. 2b,c, Suppl Figs. S2**).

### Benchmarking adaptive smoothing

We evaluated the robustness of our adaptive smoothing technique by benchmarking ClustSIGNAL against clusters generated after applying other non-spatial smoothing scenarios, such as no smoothing (Smooth0) and no smoothing with known number of clusters (Smooth0_k), and spatial smoothing scenarios, i.e., complete smoothing (Smooth100) (**Suppl Figs. S3**). For no smoothing scenario, we used the initial cluster labels generated in the first step of ClustSIGNAL as Smooth0 clustering labels. For Smooth0_k scenario, we performed a k-means clustering on the normalized data using *KmeansParam()* function in bluster package, with cluster number set to the number of clusters generated by ClustSIGNAL. In complete smoothing scenario, we used ClustSIGNAL functions to identify *N* nearest neighbours and then averaged the gene expression across all the neighbours without applying any weights. The clusters generated from this completely smoothed data were referred to as Smooth100 clusters. We ran 20 iterations of each scenario and calculated ARI and variation in number of clusters generated as measures of clustering performance (**Fig. 2d**).

### Stress testing effect of sparsity

To test if our adaptive smoothing approach was able to overcome high levels of data sparsity, we applied ClustSIGNAL to increasingly sparse data and compared its performance to that of Smooth0, Smooth0_k, and Smooth100 smoothing scenarios. For all cells in the four simulated datasets, we first downsampled the raw counts by a sparsity proportion (0, 0.1, 0.2, 0.3, 0.4, 0.5, 0.6, 0.7, 0.8, 0.9) using the *downsampleMatrix()* function in scuttle R package [68], and saved the sparse raw counts as additional as-says in the simulated data *spe* objects. Then we log-normalised each sparse assay in the four simulated data (Patched, Gradient, Complex, Uniform) using *logNormCounts()* function in scater package. We applied ClustSIGNAL, Smooth0, Smooth0_k, and Smooth100 to these assays in each dataset, with 20 iterations per test for robustness. The ability of ClustSIGNAL to cope with data sparsity was assessed based on the clustering accuracy measured as ARI (**Fig. 2e**).

### Stress testing effect of segmentation errors

We also assessed the impact of segmentation errors on the output of ClustSIGNAL and compared its performance to other smoothing scenarios (Smooth0, Smooth0_k, Smooth100). Using the *findKNN()* function from the BiocNeighbors package to identify neighbours and *downsampleMatrix()* function in scuttle package to adjust raw counts, we created various levels of segmentation errors in our simulated datasets (Patched, Gradient, Complex, Uniform) and stored the error-embedded raw counts as additional assays in the simulated data *spe* objects. To add segmentation errors to a cell *c*, we identified its five nearest neighbours and randomly selected one neighbour as the donor *d*. Following this, we first downsampled the raw counts **c** of cell *c* by a segmentation error proportion (*p_s_*), and downsampled the raw counts **d** of the donor neighbour *d* by a scaled error proportion:

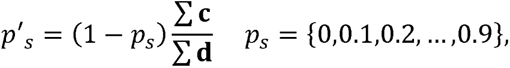

where **c** is the measured expression of cell *c*, **d** is the measured expression of donor neighbour *d*, *p_s_* is the segmentation error proportion applied to cell *c*, and *p* is the scaled error proportion applied to the neighbouring cell *d*. Then, we summed the downsampled counts from the cell *c* and its neighbour *d* to generate error-embedded raw counts for the cell *c*. For testing and comparing ClustSIGNAL, we log-normalized the error-embedded raw counts in the four simulated datasets (Patched, Gradient, Complex, Uniform) using *logNormCounts()* function in scater package, and then applied ClustSIGNAL, Smooth0, Smooth0_k, and Smooth100 to them, with 20 iterations per test for robustness. The effectiveness of ClustSIGNAL’s approach was assessed through clustering concordance measured as ARI (**Fig. 2f**).

### Stress testing reliance on initial clusters

We investigated the impact of change in initial cluster and initial subcluster labels on ClustSIGNAL’s clustering performance. For this, we assessed the effect of applying different initial clustering and sub-clustering methods, such as centroid-based clustering (minibatch k-means or mbkmeans [54]), graph-based clustering (Walktrap community detection), and density-based clustering (DBSCAN [52]), within ClustSIGNAL on the four simulated datasets (Patched, Gradient, Complex, Uniform). We performed a step-by-step ClustSIGNAL analysis, where the default clustering method (Louvain) in the first step was replaced by other clustering approaches (mbkmeans, Walktrap, DBSCAN). First, we used ClustSIGNAL with the de-fault Louvain method to note the number of initial clusters and initial subclusters generated from each simulated dataset. For mbkmeans, we used the mbkmeans R package [54] with cluster and sub-cluster numbers equal to those generated by default ClustSIGNAL. For Walktrap, we used the *buildSNNGraph()* function in scran R package to build nearest neighbour graph and *cluster_walktrap()* function in igraph R package [77] to perform community detection and generate initial clusters and initial subclusters. For DBSCAN, we used the *DbscanParam()* function in bluster package to generate clusters and sub-clusters; *NA* cluster labels were replaced by “999” to avoid errors during sub-clustering. The cluster and sub-cluster labels from all methods were then used to perform the remaining steps of ClustSIGNAL and generate final clusters. The stability of ClustSIGNAL outcome was compared in three ways: (i) by calculating the concordance of initial clusters generated by the method (mbkmeans, Walktrap, DBSCAN) compared to those generated by Louvain (default); (ii) by assessing the correlation between the entropy values generated from the method (mbkmeans, Walktrap, DBSCAN) initial subclusters *vs* those generated from Louvain (default) initial subclusters; and (iii) by calculating the concordance of final clusters generated by the method (mbkmeans, Walktrap, DBSCAN) compared to the ground truth (**Fig. 2g,h**).

### Benchmarking with existing methods

#### Spatial clustering methods

We compared ClustSIGNAL to six existing methods that perform spatial clustering. For benchmarking evaluation, we used default values for ClustSIGNAL parameters. We mainly used default or recommended parameter values, unless specified below. Method runs exceeding 400 GB RAM or 96 hours runtime were stopped due to limited availability of computational resources.

**BANKSY** is a spatial clustering method that can identify domains or cell types depending on the amount of neighbourhood representation included in the expression matrix [44]. It uses SpatialExperiment object as input, and requires pre-processing to perform joint clustering of multiple samples. We followed the recommended pre-processing steps, such as staggering spatial coordinates to ensure there was no overlap in the physical space of different samples. For all datasets, the expression counts were normalised using *NormalizeData()* function from Seurat R package [78] in accordance with BANKSY approach, and default values were used for most parameters. We used recommended values for neighbourhood size *k_geom* = *c*(15, 30), computing azimuthal Gabor filter *compute_ agf* = *TRUE* and using it *use_ agf* = *TRUE* to measure spatial pattern of gene expression, and mixing parameter *lambda* = 0.2 for cell-type clustering. For application to whole-transcriptome datasets (OC and COAD), we used 3000 highly-variable genes (as recommended in the tutorial) obtained from applying *modelGeneVar()* and *getTopHVGs()* functions from scran R package. We used BANKSY version 1.3.0 downloaded from Bioconductor: doi.org/doi:10.18129/B9.bioc.Banksy.

**BASS** is a Bayesian hierarchical model-based unsupervised method for domain and cell-type spatial clustering of SRT data [34]. It performs multi-sample clustering by taking lists of raw count matrices and spatial coordinate matrices from different samples as input. BASS requires number of cell types and do-mains as necessary inputs when creating the BASS object. Its outputs include cell-type labels, domain labels, and cell-type proportions in each spatial domain. We used default values for most parameters, except the expected number of domains and cell types. Number of cell types were selected based on prior annotations, while domain numbers were either randomly selected or were taken from published data: 23 cell types and 10 domains (random selection) for ME; 19 cell types and 7 domains (Fig. 4d in Janesick *et al* [48]) for BR; 22 cell types and 9 domains (niches defined in He *et al* [9]) for LC; 16 cell types and 9 domains (Fig. 3e in Moffitt *et al* [47]) for MH; 13 cell types and 5 domains (niches defined in Ren *et al* [49]) for OC; 16 cell types and 8 domains (niches defined in Ren *et al* [49]) for COAD. For benchmarking, we used the cell-type labels to assess method performance. By default, BASS internally selects 3000 spatially-variable genes for clustering, and would have applied this limit to whole-transcriptome datasets (OC and COAD). We used BASS version 1.1.0.017 downloaded from GitHub: github.com/zhengli09/BASS, with modifications to its C library version (changed from CXX11 to CXX14) to avoid dependency issues. Furthermore, to run BASS on datasets with large samples, such as BR, LC, OC, and COAD, we had to increase the C stack memory size of the run environment from 8 to 64 MB.

**SpatialPCA** is a spatially-aware dimension reduction method that expands on the probabilistic model of PCA [45]. While SpatialPCA comes with a multi-sample analysis script, we incurred errors when we tried running it on a real-world dataset. Therefore, we performed SpatialPCA clustering on individual samples, and the cluster labels do not match between the samples. We followed the method tutorial and used default or recommended parameter values for large datasets, setting *sparkversion* = *sparkx* for selecting spatial genes, *sparseKernel* = *TRUE* to use sparse matrix kernel, and *bandwidthtype* = *Silverman* for Gaussian kernel. Furthermore, to ensure SpatialPCA would run on large datasets, we increased the threshold for building the sparse kernel matrix *sparseKernel_tol* = 1*e^−^*^5^, i.e., any values below this cut-off would be set to zero in the matrix. By default, SpatialPCA internally selects up to 3000 significantly spatially-variable genes for dimension reduction (*gene.number* = 3000, *gene.type* = “spatial”), and would have applied this limit to whole-transcriptome datasets (OC and COAD). We used SpatialPCA version 1.3.0 downloaded from GitHub: github.com/shangll123/SpatialPCA.

**SpaGCN** is a DL-based spatial clustering method that integrates information from gene expression, spatial locations, and optionally histology features using a graph convolutional network, followed by iterative clustering [30]. It requires pre-processing, such as staggering spatial coordinates of samples, to perform joint clustering of multiple samples. We used default and recommended values to run SpaGCN on each sample of the six datasets, and set the number of clusters (*n_clusters*) equal to the number of cell-type annotations in each dataset: 23 for ME, 19 for BR, 22 for LC, 16 for MH, 13 for OC, and 16 for COAD. We did not use histology information as this information was not readily available for the selected datasets. SpaGCN exceeded memory limit during spatial graph generation for OC and COAD, as these datasets contain single samples with a large number of cells, and applying any gene selection to these datasets would not be helpful. We downloaded SpaGCN v1.2.7 from GitHub: github.com/jianhuupenn/SpaGCN and ran it on a Python 3.13.5 environment with relevant dependencies installed as instructed in tutorials.

**STAGATE** is also a DL-based spatial clustering method [33]. It uses an attention mechanism to adaptively aggregate neighbourhood expression into a low dimensional embedding, followed by clustering. We mainly used default and recommended values for benchmarking the six methods. However, due to the large differences between the spatial coordinate systems of high-resolution technologies, we used the k-nearest neighbour approach with *k* = 30 (same as ClustSIGNAL) to select neighbourhoods, in place of the default radius threshold approach. For application to whole-transcriptome datasets (OC and COAD), we used 3000 highly-variable genes obtained from applying *sc.pp.highly_variable_genes()* function from scanpy Python package, as per instructions in the tutorial. For the final clustering step, we replaced the recommended mclust model-based clustering with the Louvain community detection method due to issues with running the mclust R package in a Python environment. We downloaded STAGATE_pyG from GitHub: github.com/RucDongLab/STAGATE_pyG and ran it on a Python 3.13.5 environment with relevant dependencies installed as instructed in tutorials.

**GraphST** is another DL-based method that relies on contrastive learning to generate a refined embed-ding of gene expression and spatial information and perform spatial clustering [36]. It requires alignment of samples prior to multi-sample analysis. We used default values for running GraphST on each sample of the six datasets, and set the number of clusters (n clusters) equal to the number of cell-type annotations in each dataset, same as for SpaGCN. However, for the final clustering step, we replaced the default mclust clustering method with the Louvain community detection method (*method* = “louvain”), same as for STAGATE. GraphST was only able to run on the smallest ME dataset containing 50,000 cells within a 400 GB memory and 96 hours runtime limit. We downloaded GraphST v1.1.1 from GitHub: github.com/JinmiaoChenLab/GraphST and ran it on a Python 3.8.19 environment with relevant dependencies installed as instructed in tutorials.

#### Performance evaluation

We assessed the performance of the methods based on their clustering concordance measured in terms of ARI and their computational efficiency measured by runtime and memory usage on six real-world SRT datasets. For each dataset, ARI was calculated between prior annotations and method clusters using *ARI()* function from aricode R package. Memory usage and runtime were measured using the *peakRAM()* function in peakRAM R package, as peak RAM used and elapsed time, respectively. All methods, including DL methods, were run on a single CPU core for fair comparison between methods.

### Comparison of integration and multi-sample analysis

We developed ClustSIGNAL for multi-sample analysis of high-resolution SRT datasets. For this, ClustSIGNAL takes a single, combined SpatialExperiment object of all samples as input, with sample names as additional information when running the method. During the neighbourhood detection and sorting steps, ClustSIGNAL searches for neighbours in individual samples followed by compilation of entropies from all samples into a singular framework (**Suppl Fig. S27**). For datasets with strong batch effects, we added an optional functionality to call Harmony [69] for batch integration. To assess ClustSIGNAL’s multi-sample analysis approach, we performed the following comparisons: (i) ClustSIGNAL *vs* PASTE + ClustSIGNAL, where we compared ClustSIGNAL with and without PASTE [79] spatial alignment of 3 mouse embryo (ME) samples [46] to investigate the impact of spatial integration on ClustSIGNAL’s performance; (ii) ClustSIGNAL *vs* GraphST, where we compared ClustSIGNAL with spatial alignment of ME samples against GraphST [36], which was designed for spatial integration of samples to perform multi-sample analysis; and (iii) ClustSIGNAL *vs* Harmony + ClustSIGNAL, where we compared ClustSIGNAL performance with and without Harmony integration of lung cancer (LC) data [9] with strong batch effects. We assessed the clustering performance using ARI as a measure of concordance between clusters and prior annotations.

### Marker identification and enrichment analyses

We assessed the biological relevance of specific clusters by identifying marker genes in the clusters and performing gene ontology (GO) over-representation analysis. For comparison between specific clusters, the dataset was first subset to include only the cells belonging to the clusters under investigation. We applied the *findMarkers()* function from scran R package and a pairwise t-test to cells grouped by cluster labels to identify differentially expressed genes. Then, we selected the top 10 significantly differential genes (Benjamini-Hochberg (BH)-adjusted *p*-value (FDR) *<* 0.05) from each cluster as markers. In seqFISH mouse embryo dataset, the “Xist” gene associated with X-chromosome activation was excluded before analysing the clusters. To assess the biological processes associated with the marker genes, we performed a GO over-representation analysis. For this, we first selected all genes that were significantly upregulated in the cluster (*FDR <* 0.05; *logFC >* 0.2). We applied the *enrichGO()* function from clusterProfiler R package [80] to these gene names using the org.Hs.eg.db R package for human gene annotation database, org.Mm.eg.db R package for mouse gene annotation database, *BH* method for *p*-value adjustment, and significant unadjusted (*pvalueCutoff* = 0.05) and adjusted (*qvalueCutoff* = 0.05) *p*-values. We specifically searched through biological processes (BP) ontologies, and mainly considered processes associated with at least 5 genes.

### Co-localisation and cell type neighbourhood analyses

We studied the co-localisation of cluster cells using the neighbourhood data generated during ClustSIGNAL runs on the real-world datasets. In a sample, we used each cell and their 30-nearest neighbours to generate a cluster-by-cluster matrix, where the rows were cluster labels of index cells (index clusters), columns were cluster labels of neighbouring cells (neighbouring clusters), and the values indicated the number of neighbourhood cells that belonged to specific neighbouring clusters. We further normalised each row by the total number of cells in the row (i.e., total cells in the index cluster), giving us a matrix of cell number proportions in a sample. Similarly, for cell type neighbourhood analysis, we used the neighbourhood data generated from application of ClustSIGNAL to MH dataset. We assessed the association of Inhibitory and Excitatory neuron cluster cells with the prior annotations by calculating whether a cell type was detected in the neighbourhood (30-nearest neighbours) of a cluster cell. We further assessed the statistical significance of these associations using a chi-square test with BH correction.

### Pseudotime trajectory analysis

We performed a trajectory analysis on the cells from the ME data to assess whether ClustSIGNAL clusters represented biologically meaningful cell states along a developmental lineage. For this, we applied the *slingshot()* function from the slingshot R package [81] to the top 20 PCs generated from the normalised gene expression of all 3 embryos, and used ClustSIGNAL clusters for the cluster-based minimum spanning tree. For visualisation, we additionally applied the *embedCurves()* function to project the smooth trajectory curves onto the UMAP space.

#### Spatial proteomics data entropy analysis

We evaluated the entropy values generated by ClustSIGNAL to explore their use in data analysis and gaining biological insights from disease samples. For this, we used a publicly-available spatial proteomics dataset of breast cancer tissues from Keren *et al* [82] to identify any association between entropy values and patient survival. The dataset contained intensities of 48 features in 197,678 cells from 40 TNBC (triple negative breast cancer) patient samples, of which we removed 11 features that represented metal ions (Na, Si, P, Ca, Fe, Ta, Au), control (Background), and nuclear stains (dsDNA, H3K9ac, H3K27me3). We calculated expression values from intensities using the protocol reported in Keren *et al* [82], i.e., dividing the feature intensities in each cell by cell sizes, followed by Arcsinh transformation and z-score normalisation of the data. We used these normalised expression values to calculate PCA and UMAP low embeddings using *runPCA()* and *runUMAP()* functions, respectively, in scater package and identify any batch effects. We applied ClustSIGNAL to the normalised expression to generate entropy values, performing batch correction by Group. For comparison between samples, we normalised the entropy values (*E_c_*) in each sample using the log of total initial subcluster labels (*K*) in a sample:

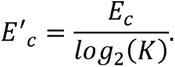

We further assessed the correlation between survival time and normalised entropy-based metrics per sample, such as mean, median, and standard deviation of entropy as well as proportion of high (*entropy >* 0.6) and low (*entropy <* 0.4) entropy values, focussing on the 15 samples for which uncensored survival data were available (see **Supplementary Material**).

## Data availability

The six publicly-available SRT datasets we used for benchmarking and real data analysis were: (i) SeqFISH mouse embryo dataset downloaded from Spatial Mouse Atlas website, (ii) Xenium breast cancer dataset downloaded from 10x Genomics website and annotations acquired from source data available from Janesick *et al* [48], (iii) CosMx lung cancer data taken from Nanostring website, (iv) MERFISH mouse hypothalamus preoptic region dataset downloaded from the DRYAD website, (v) Stereo-seq ovarian cancer dataset downloaded from the SPATCH website, and (vi) VisiumHD colon cancer dataset downloaded from the SPATCH website. For simulated data analysis, we used the Tabula Sapiens human tissue scRNA-seq atlas hosted on the CellxGene website. For exploration of entropy, we used a breast cancer spatial proteomics data accessed from SpatialDatasets R package.

## Code availability

ClustSIGNAL R package is available on Bioconductor from bioconductor.org/packages/clustSIGNAL and the latest version with faster implementation of ClustSIGNAL (v1.5.2) is available on GitHub from SydneyBioX/clustSIGNAL. The scripts for simulation, benchmarking, and data analysis are available on GitHub from SydneyBioX/ClustSIGNAL analysis2025.

## Acknowledgements

We thank our colleagues in the University of Sydney School of Mathematics and Statistics, Charles Perkins Centre, and Sydney Precision Data Science Centre for their support and intellectual engagement through constructive comments and suggestions.

## Author Contributions

S.G. and S.C.H. conceived the study. P.P. conceptualized the methodological framework of ClustSIGNAL with suggestions from S.G., B.G., H.Z., and S.C.H.. P.P. led the development of the method and software package. P.P performed the formal analyses and interpretation of results under the guidance of S.G.. P.P. and H.Z. performed benchmarking analysis under the guidance of S.G., B.G., and S.C.H.. P.P. wrote the initial draft of the manuscript, and all authors read, edited, and approved the final version of the manuscript.

