## Supplementary Material for "ClustSIGNAL identifies cell types and subtypes using an adaptive smoothing approach for scalable spatial omics clustering"

#### Assessing ClustSIGNAL's multi-sample approach

We evaluated the effect of spatial alignment of mouse embryo (ME) [1] samples using PASTE [2] on the performance of ClustSIGNAL, and found negligible differences in cluster concordance with prior annotations (**Suppl Fig. S28**). This suggested that spatial alignment does not impact ClustSIGNAL's performance, which is supported by ClustSIGNAL's approach of identifying neighbours in individual samples followed by compilation of entropy values from all samples (**Suppl Fig. 27**). We then compared ClustSIGNAL's performance on the ME data to that of GraphST, a spatial clustering method with inherent batch correction capabilities [3]. ClustSIGNAL (mean ARI: 0.49) performed better clustering than GraphST with (mean ARI: 0.38) or without (mean ARI: 0.24) PASTE alignment of the samples (**Suppl Fig. S28**). This also highlighted that spatial alignment of samples is not always beneficial when performing multi-sample analysis, as GraphST performance was worse on aligned ME samples compared to per-sample analysis. Finally, we compared ClustSIGNAL's performance with and without using Harmony [4] on 8 lung cancer (LC) tissue slices [5] with strong batch effects. ClustSIGNAL performance improved with application of Harmony, showing higher and less variable (mean ARI $\pm$ SD: 0.47 $\pm$ 0.2 *vs* 0.53 $\pm$ 0.1) cluster concordance with prior annotations (**Suppl Fig. 27**).

#### Exploring ambiguous clusters in breast cancer and lung cancer data

In the Xenium breast cancer (BR) [6] and CosMx lung cancer (LC) [5] datasets, we noticed that ClustSIGNAL often partitioned ambiguous cells into distinct clusters (**Suppl Figs. S32, S33**). In the BR data, the Invasive_Tumor cells were sub-clustered into 3 different clusters (2, 3, 9), of which clusters 3 and 9 represented subgroups of the Invasive_Tumor region, whereas cluster 2 cells occupied the area around the edges of the Invasive_Tumor, closer to the tumour micro-environment (**Suppl Figs. S32a-c**). Further analysis of cluster 2 showed that it contained both tumour and immune marker genes, and likely represented cells with segmentation errors (**Suppl Figs. S32d,e,i**).

In the LC data, we observed that the clusters obtained from ClustSIGNAL, after Harmony integration of the samples by patient group, contained subgroups of prior annotations. Notably, tumor 9 annotated cells from two Lung9 samples were split into two clusters (7, 10), one cluster per sample (**Suppl Figs. S33a,b**). Analysis of the marker gene expression and cluster co-localisation revealed that these two groups of cells had distinct transcriptomics profiles (**Suppl Figs. S33c-e**). We also observed intra-slice batch effects in some field of views (FOVs) of Lung6 sample (**Suppl Figs. S33f**). While most tumour cells in this sample were grouped under cluster 8, the cells belonging to certain FOVs were grouped under cluster 12. Notably, these prominent FOV batch effects were not observed when ClustSIGNAL was applied to the LC data without Harmony integration (**Suppl Figs. S6**). This highlights ClustSIGNAL's inherent robustness to spatial artifacts such as FOV batch effects.

#### Exploring association between entropy and patient survival time in spatial proteomics data

A by-product of applying ClustSIGNAL to any dataset is the generation of cell-specific entropy values that measure the heterogeneity level of a cell's neighbourhood. We assessed the use of entropy as an additional metric for data exploration and gaining biological insights, beyond clustering. For this, we applied ClustSIGNAL to a spatial proteomics breast cancer dataset [7] containing 40 samples from different patients, and analysed the relationship between entropy and patient survival using only 15 samples for which uncensored survival data were available in the dataset (**Suppl Fig. 34**). For fair comparison between samples, we first normalised the entropy values and then generated sample-specific entropy metrics (**Suppl Fig. 34b,c**; also see **Methods**). We observed association between patient survival time and mean entropy, median entropy, and proportion of high or low entropy. Of these, the proportion of high and low entropy regions showed the strongest associations with patient survival, with patient tissues that had a larger proportion of low entropy regions, i.e., homogeneous neighbourhoods with less mixing of different cell types, being linked to a lower survival time, whereas patient tissues with larger proportion of high entropy regions, i.e., heterogeneous neigbourhoods with a diverse population of cell types, being associated with a higher survival time (**Suppl Fig. 34c**).

### Supplementary Tables

**Suppl Table S1:** Benchmarking results from applying ClustSIGNAL and six other spatial clustering methods to six spatially-resolved transcriptomics datasets from different technologies. *For benchmarking with Stereo-seq and VisiumHD datasets, top 3000 highly-variable genes were used by the methods. †Running BASS on datasets with large samples, such as BR, LC, OC, and COAD, required extending R environment's C stack memory size from 8 MB (default) to 64 MB. NA (not available) indicates method runs that failed, either due to excessive memory usage (>400 GB) or excessive runtime (>96 hours).

| Dataset | Samples | Cells | Genes | Metric | ClustSIGNAL | BANKSY | BASS† | SpatialPCA | SpaGCN | STAGATE | GraphST |
|---|---|---|---|---|---|---|---|---|---|---|---|
| SeqFISH mouse embryo | 3 | 52,568 | 351 | Mean ARI | **0.49** | 0.45 | 0.45 | 0.26 | 0.44 | 0.16 | 0.38 |
| | | | | Runtime (hr) | **0.03** | 0.03 | 1.2 | 10.85 | 0.29 | 3.79 | 3.25 |
| | | | | Peak RAM (GB) | **0.91** | 2.9 | 16.75 | 39.95 | 7.53 | 8.65 | 28.44 |
| Xenium breast cancer | 2 | 274,501 | 313 | Mean ARI | **0.48** | 0.35 | 0.49 | *NA* | 0.38 | 0.14 | *NA* |
| | | | | Runtime (hr) | **0.42** | 0.27 | 16.7 | *NA* | 12.83 | 20.6 | *NA* |
| | | | | Peak RAM (GB) | **3.4** | 12.95 | 103.67 | *NA* | 286.95 | 51.74 | *NA* |
| Stereo-seq ovarian cancer | 1 | 284,984 | 31,585* | Mean ARI | **0.65** | 0.55 | 0.5 | *NA* | *NA* | 0.21 | *NA* |
| | | | | Runtime (hr) | **0.81** | 0.69 | 20.21 | *NA* | *NA* | 8.13 | *NA* |
| | | | | Peak RAM (GB) | **14.1** | 72.58 | 87.62 | *NA* | *NA* | 104.37 | *NA* |
| VisiumHD colon cancer | 1 | 520,382 | 18,042* | Mean ARI | **0.38** | 0.42 | *NA* | *NA* | *NA* | 0.2 | *NA* |
| | | | | Runtime (hr) | **1.55** | 2.77 | *NA* | *NA* | *NA* | 53.23 | *NA* |
| | | | | Peak RAM (GB) | **15.05** | 131.42 | *NA* | *NA* | *NA* | 181.8 | *NA* |
| CosMx SMI lung cancer | 9 | 771,236 | 960 | Mean ARI | **0.47** | 0.5 | 0.55 | *NA* | 0.33 | 0.07 | *NA* |
| | | | | Runtime (hr) | **2.08** | 2.82 | 20.44 | *NA* | 42.62 | 60.63 | *NA* |
| | | | | Peak RAM (GB) | **20.25** | 64.1 | 293.39 | *NA* | 295.86 | 53.76 | *NA* |
| MERFISH mouse hypothalamus | 181 | 874,756 | 135 | Mean ARI | **0.45** | 0.39 | 0.43 | 0.09 | 0.33 | 0.05 | *NA* |
| | | | | Runtime (hr) | **2.04** | 3.43 | 15.48 | 18.48 | 2.88 | 59.22 | *NA* |
| | | | | Peak RAM (GB) | **6.84** | 21.4 | 272.96 | 3.74 | 6.71 | 7.95 | *NA* |

**Suppl Table S2:** Composition of reference cell types used to generate cell labels in each simulation pattern. Conjunctiva, conjunctival epithelial cell; Cornea, corneal epithelial cell; Kidney, Kidney epithelial cell; Mesenchyma, mesenchymal stem cell; Retina, retinal blood vessel endothelial cell; Skeletal muscle, skeletal muscle satellite stem cell; NA, not applicable.

| Cell labels | Patched | Gradient | Complex | Uniform |
|---|---|---|---|---|
| 1 | B cell | B cell | B cell | B cell |
| 2 | B cell, Plasma cell | B cell, Plasma cell | Plasma cell | Plasma cell |
| 3 | Plasma cell | Plasma cell | Conjunctiva | Conjunctiva |
| 4 | B cell, Conjunctiva | Plasma cell, Conjunctiva | Hepatocyte | Cornea |
| 5 | Retina | Conjunctiva | Conjunctiva, Cornea | Retina |
| 6 | Plasma cell, Cornea | Cornea | Hepatocyte, Kidney | Kidney |
| 7 | Conjunctiva | Retina | Cornea | Hepatocyte |
| 8 | Conjunctiva, Cornea | Retina, Kidney | Kidney | Mesenchyma |
| 9 | Cornea | Kidney | Mesenchyma | Skeletal muscle |
| 10 | *NA* | Hepatocyte | Skeletal muscle | B cell, Plasma cell |
| 11 | *NA* | Hepatocyte, Mesenchyma | Mesenchyma, Skeletal muscle | Conjunctiva, Cornea |
| 12 | *NA* | Mesenchyma | B cell, Plasma cell | Mesenchyma, Skeletal muscle |
| 13 | *NA* | Mesenchyma, Skeletal muscle | *NA* | *NA* |
| 14 | *NA* | Skeletal muscle | *NA* | *NA* |

#### Supplementary Figures

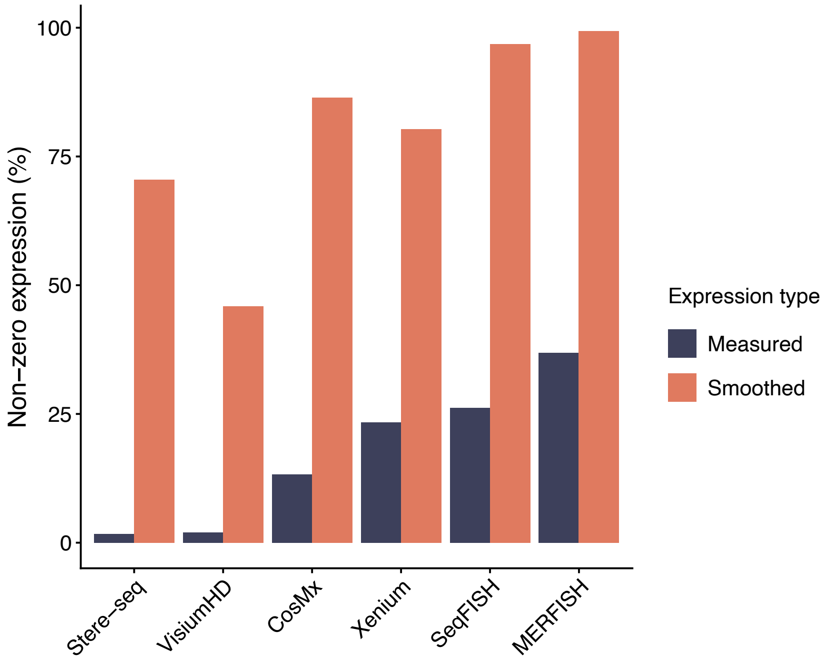


**Suppl Fig. S1:** Bar plot showing level of sparsity in spatial transcriptomics data obtained from various technologies. Dark blue bars represent the raw counts, i.e., the total transcripts, profiled using the platforms, and orange bars indicate the adaptively smoothed expression generated by ClustSIGNAL. Stereo-seq, human ovarian cancer data (OC) [8]; VisiumHD, human colon cancer data (COAD) [8]; CosMx, human lung cancer data (LC) [5]; Xenium, human breast cancer data (BR) [6]; SeqFISH, mouse embryo data (ME) [1]; MERFISH, mouse hypothalamus preoptic region data (MH) [9].

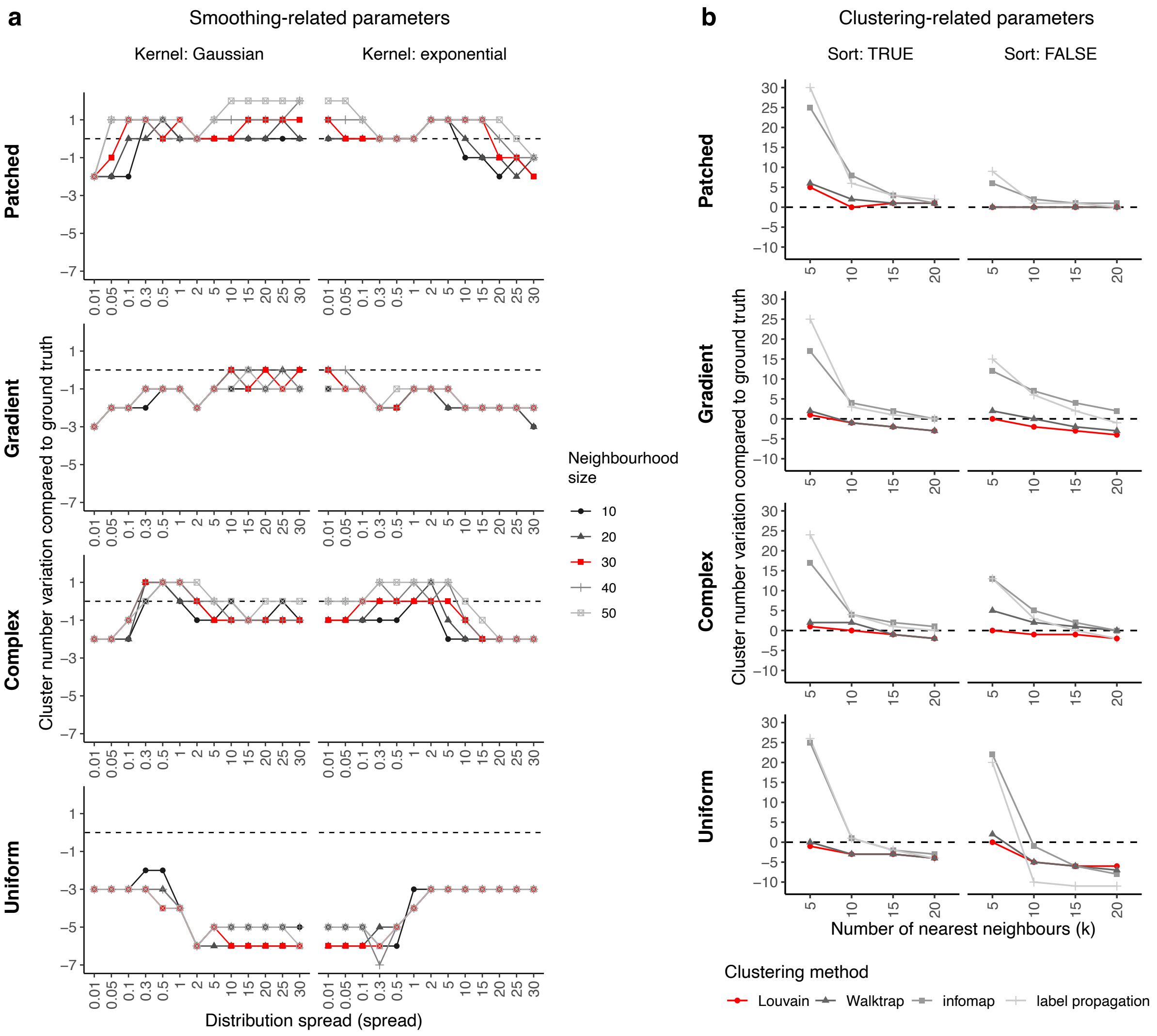


**Suppl Fig. S2:** Plots showing parameters associated with **(a)** adaptive smoothing, i.e., neighbourhood size ($N$), kernel type (Gaussian, exponential), and kernel spread (standard deviation of Gaussian kernel or rate of exponential kernel), and **(b)** clustering and other components, such as number of nearest neighbours ($k$) in graph clustering, community detection method (Louvain, Walktrap, infomap, label propagation), and neighbourhood sorting. Optimal values for the parameters were selected based on ARI **(Fig. 2b,c)** and number of clusters identified compared to ground truth **(a, b)**.

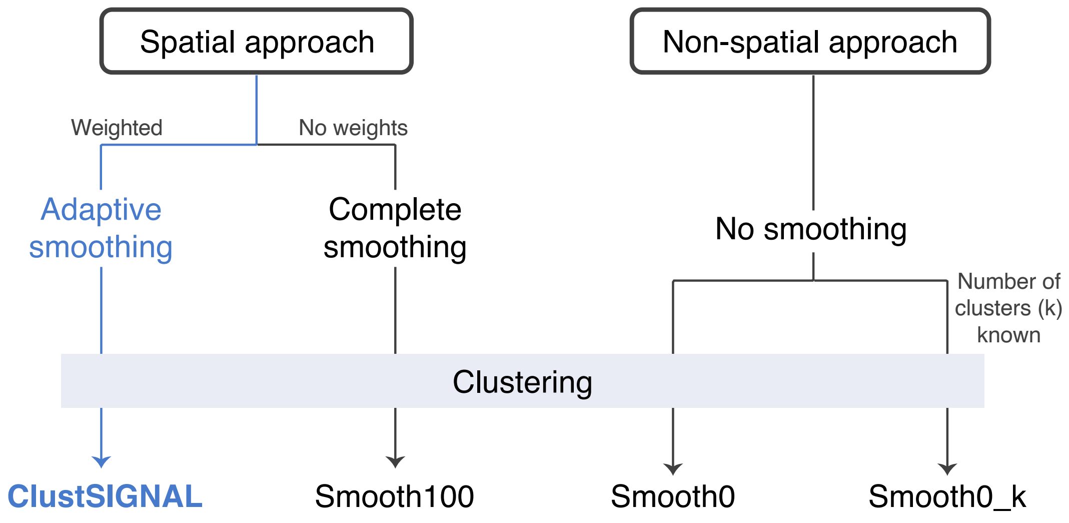


**Suppl Fig. S3:** Schematic overview of the simulated data analysis to assess the robustness of ClustSIGNAL's adaptive smoothing approach compared to other smoothing scenarios (no smoothing and complete smoothing).

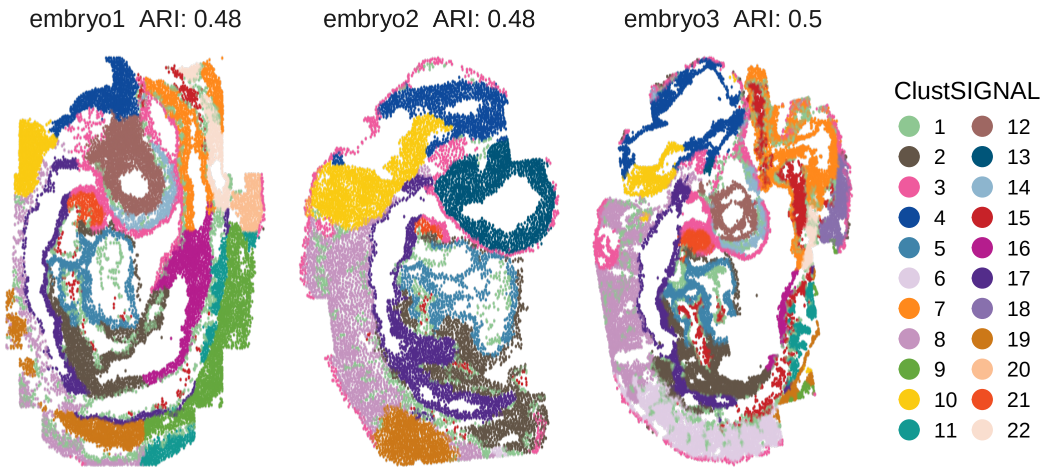


**Suppl Fig. S4:** Spatial plots of ClustSIGNAL clusters in seqFISH mouse embryo data [1]. ARI between ClustSIGNAL clusters and annotations were calculated for each sample as a measure of clustering concordance.

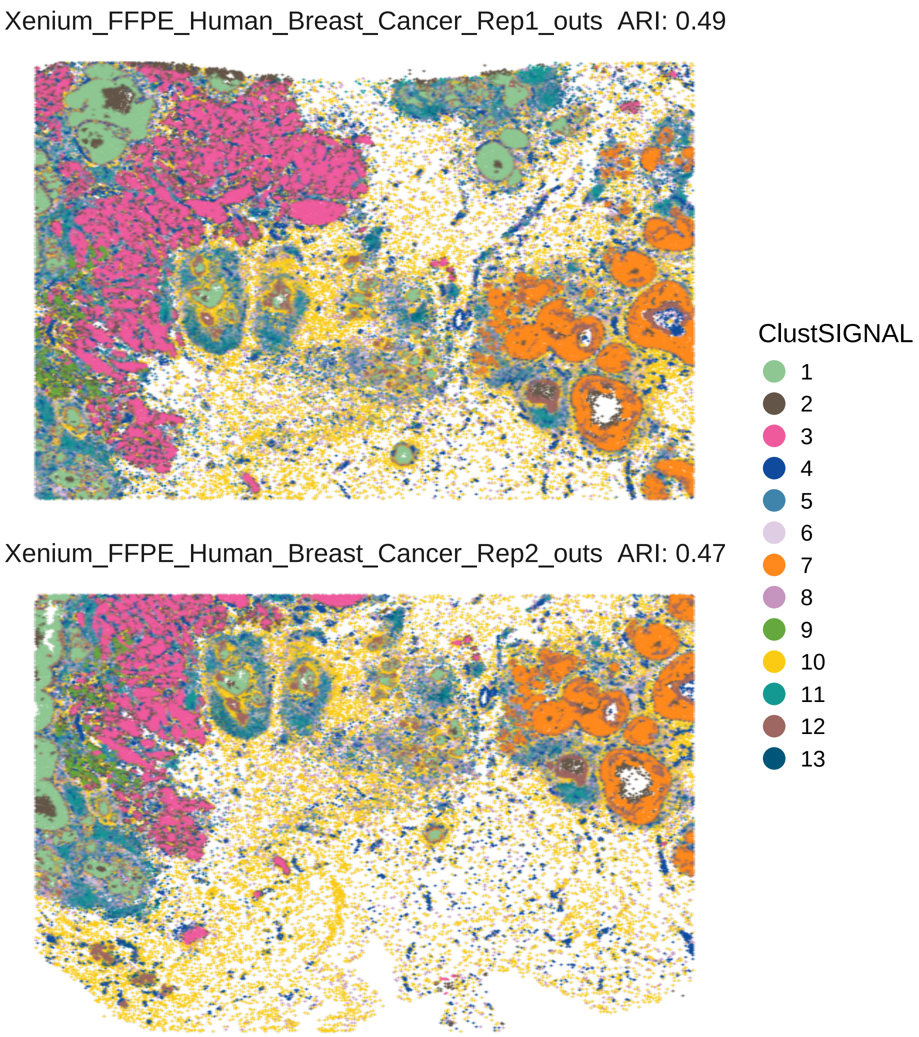


**Suppl Fig. S5:** Spatial plots of ClustSIGNAL clusters in Xenium breast cancer data [6]. ARI between ClustSIGNAL clusters and annotations were calculated for each sample as a measure of clustering concordance.

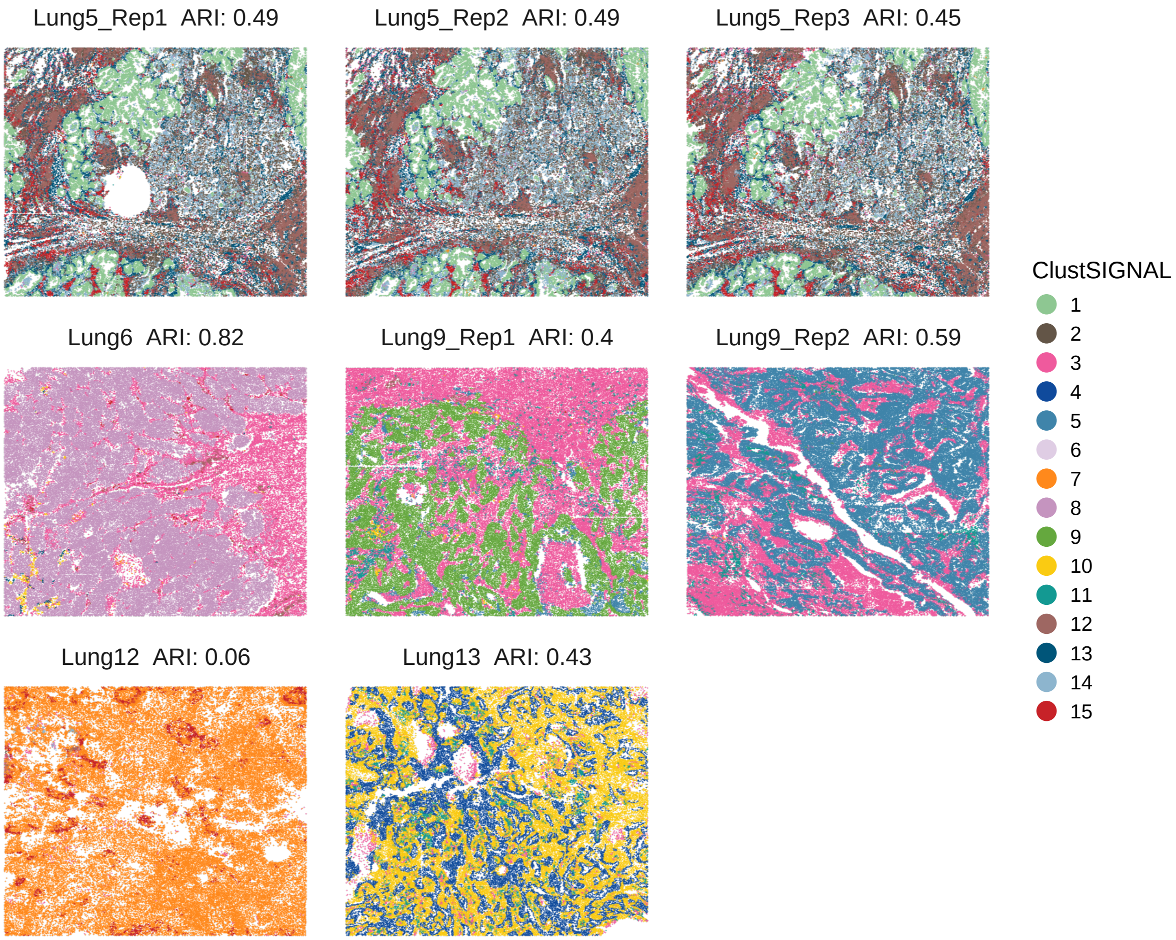


**Suppl Fig. S6:** Spatial plots of ClustSIGNAL clusters in CosMx lung cancer data [5]. ARI between ClustSIGNAL clusters and annotations were calculated for each sample as a measure of clustering concordance.

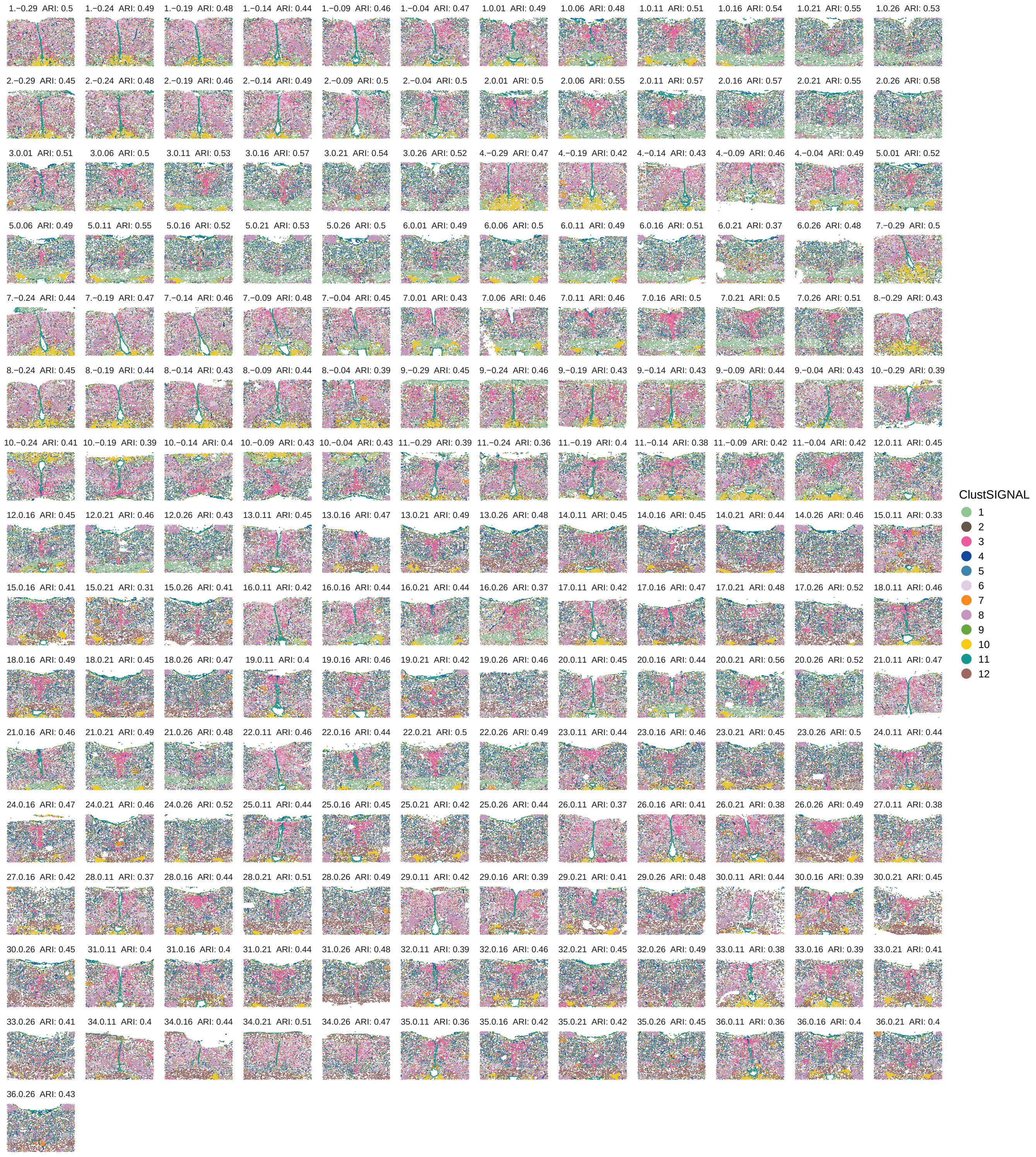


**Suppl Fig. S7:** Spatial plots of ClustSIGNAL clusters in MERFISH mouse hypothalamus preoptic region data [9]. ARI between ClustSIGNAL clusters and annotations were calculated for each sample as a measure of clustering concordance.

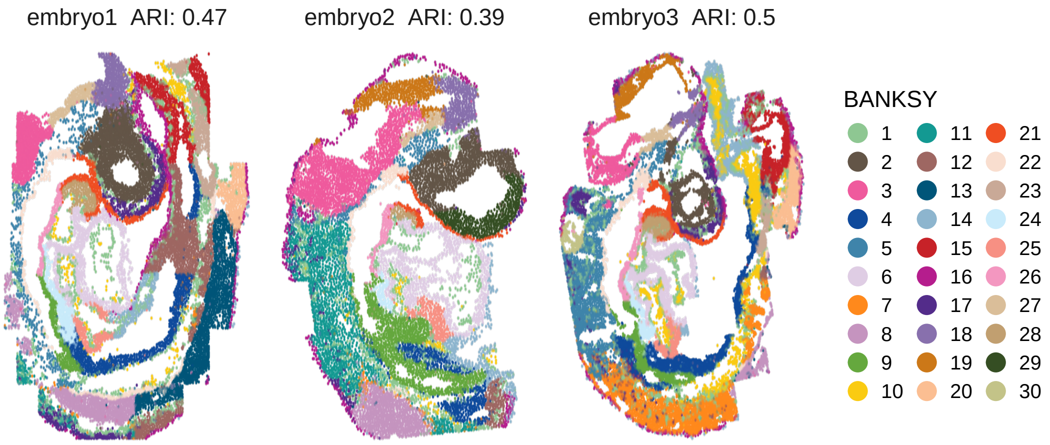


**Suppl Fig. S8:** Spatial plots of BANKSY clusters in seqFISH mouse embryo data [1]. ARI between BANKSY clusters and annotations were calculated for each sample as a measure of clustering concordance.

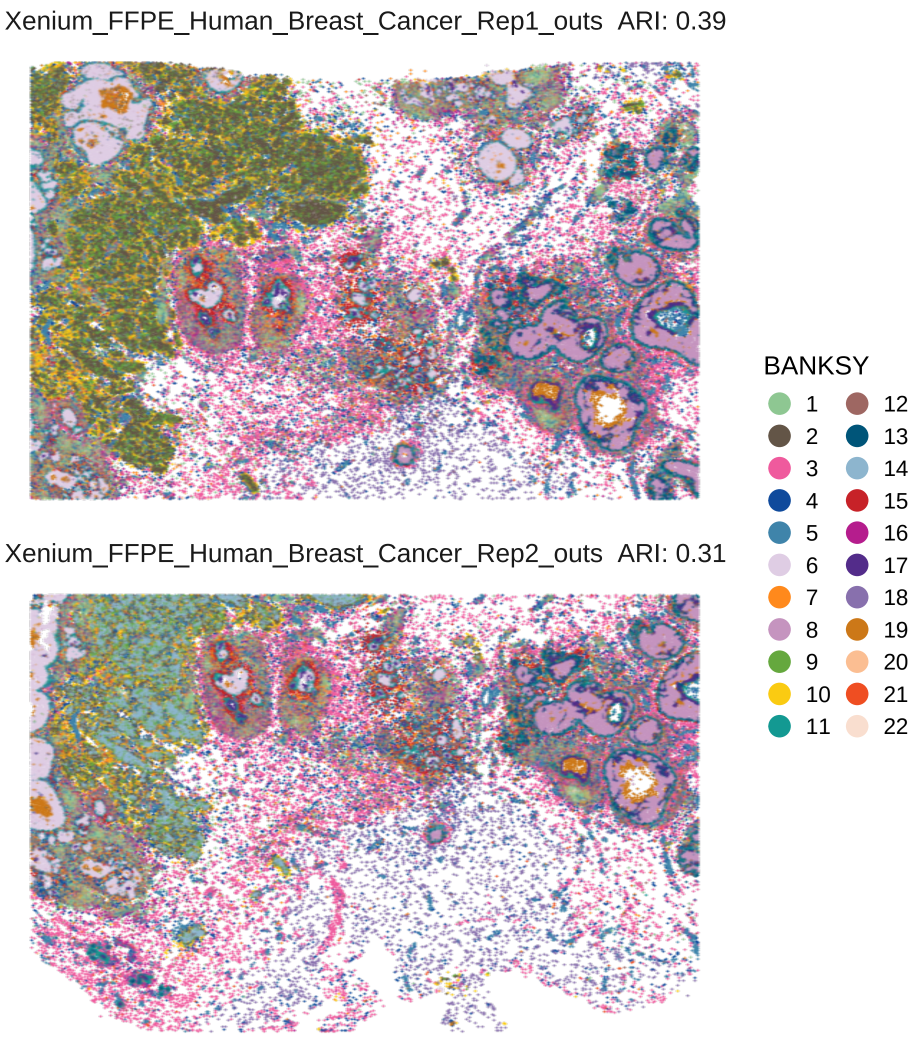


**Suppl Fig. S9:** Spatial plots of BANKSY clusters in Xenium breast cancer data [6]. ARI between BANKSY clusters and annotations were calculated for each sample as a measure of clustering concordance.

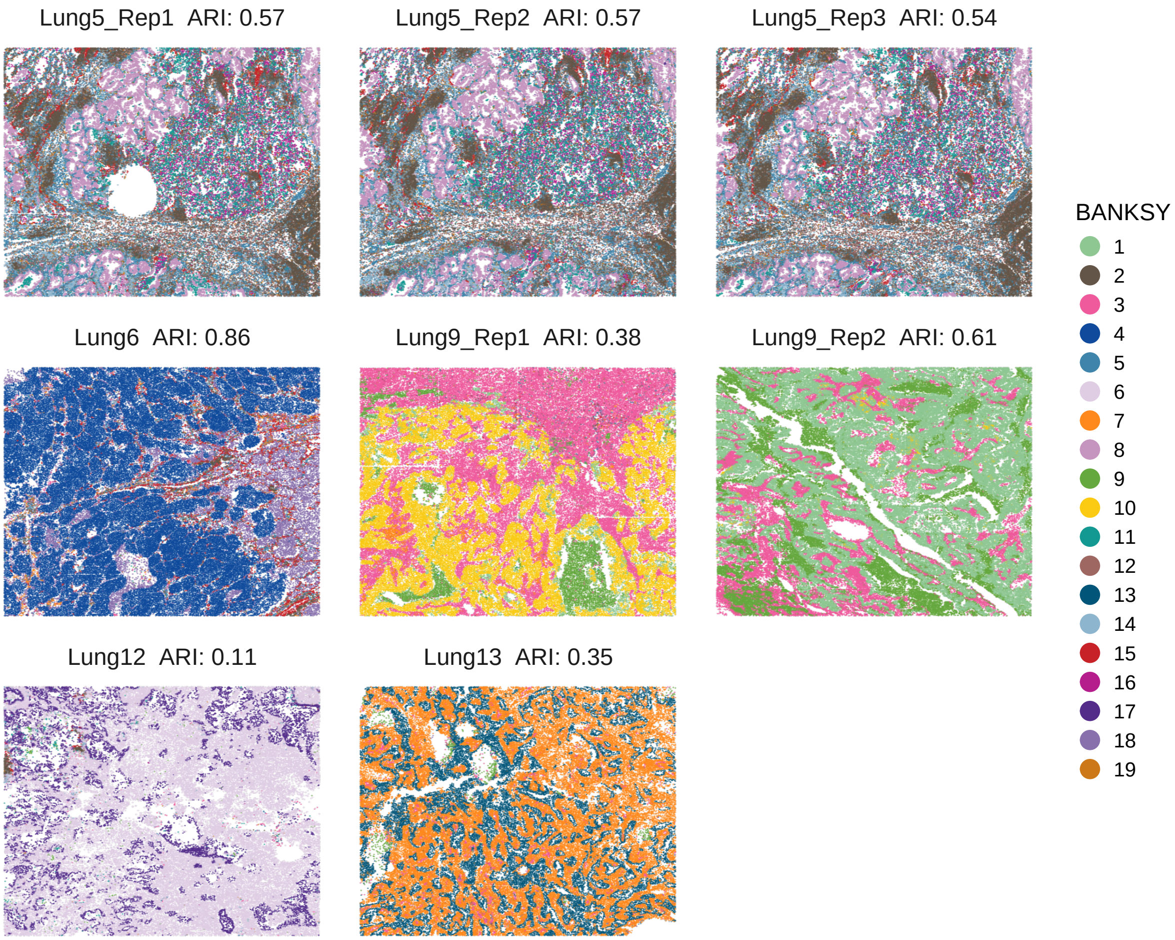


**Suppl Fig. S10:** Spatial plots of BANKSY clusters in CosMx lung cancer data [5]. ARI between BANKSY clusters and annotations were calculated for each sample as a measure of clustering concordance.

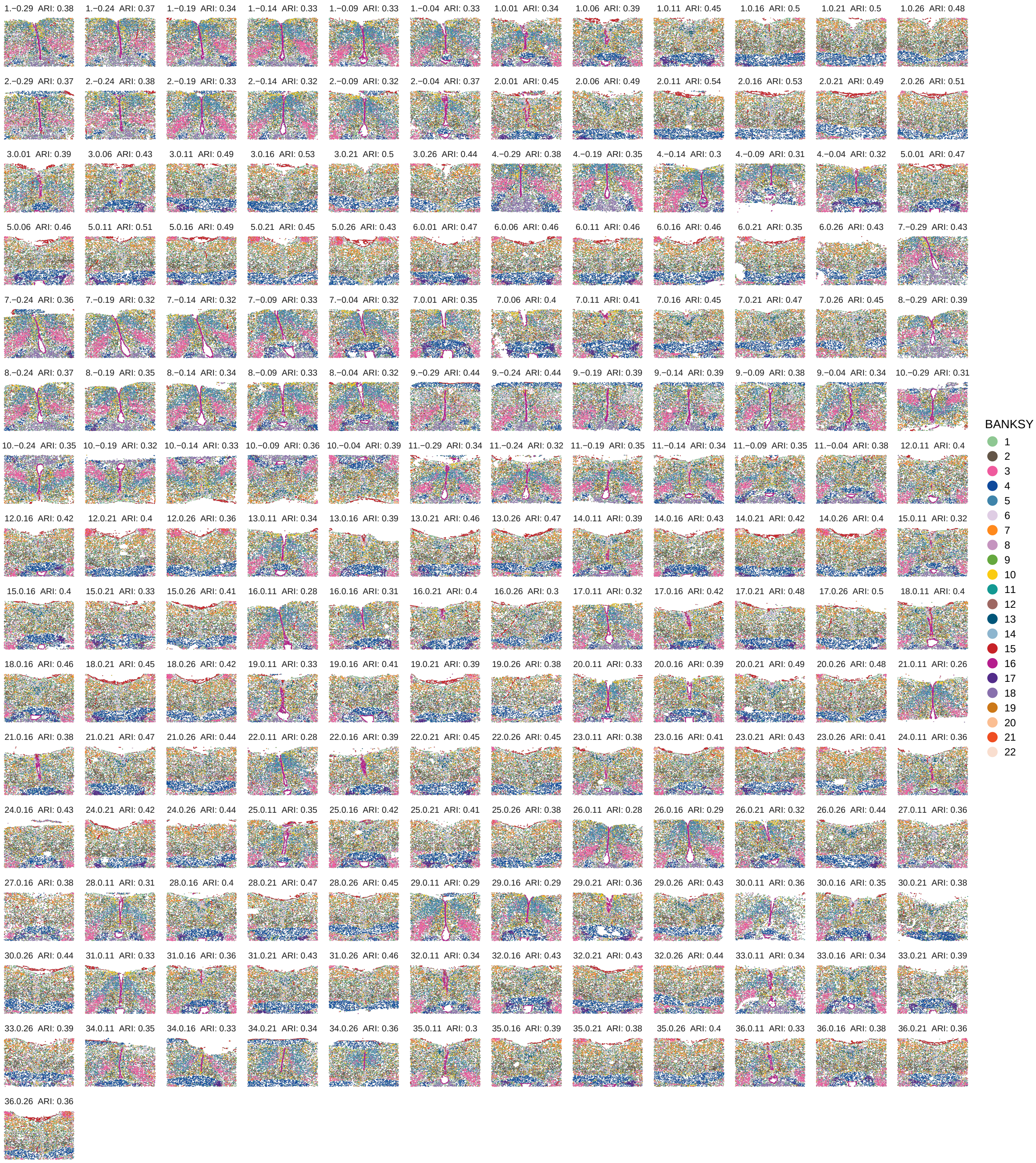


**Suppl Fig. S11:** Spatial plots of BANKSY clusters in MERFISH mouse hypothalamus preoptic region data [9]. ARI between BANKSY clusters and annotations were calculated for each sample as a measure of clustering concordance.

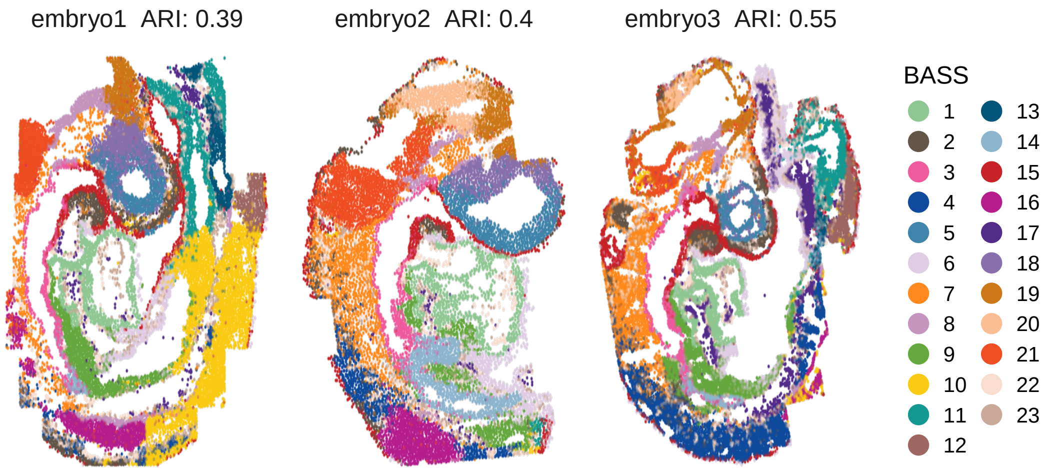


**Suppl Fig. S12:** Spatial plots of BASS clusters in seqFISH mouse embryo data [1]. ARI between BASS clusters and annotations were calculated for each sample as a measure of clustering concordance.

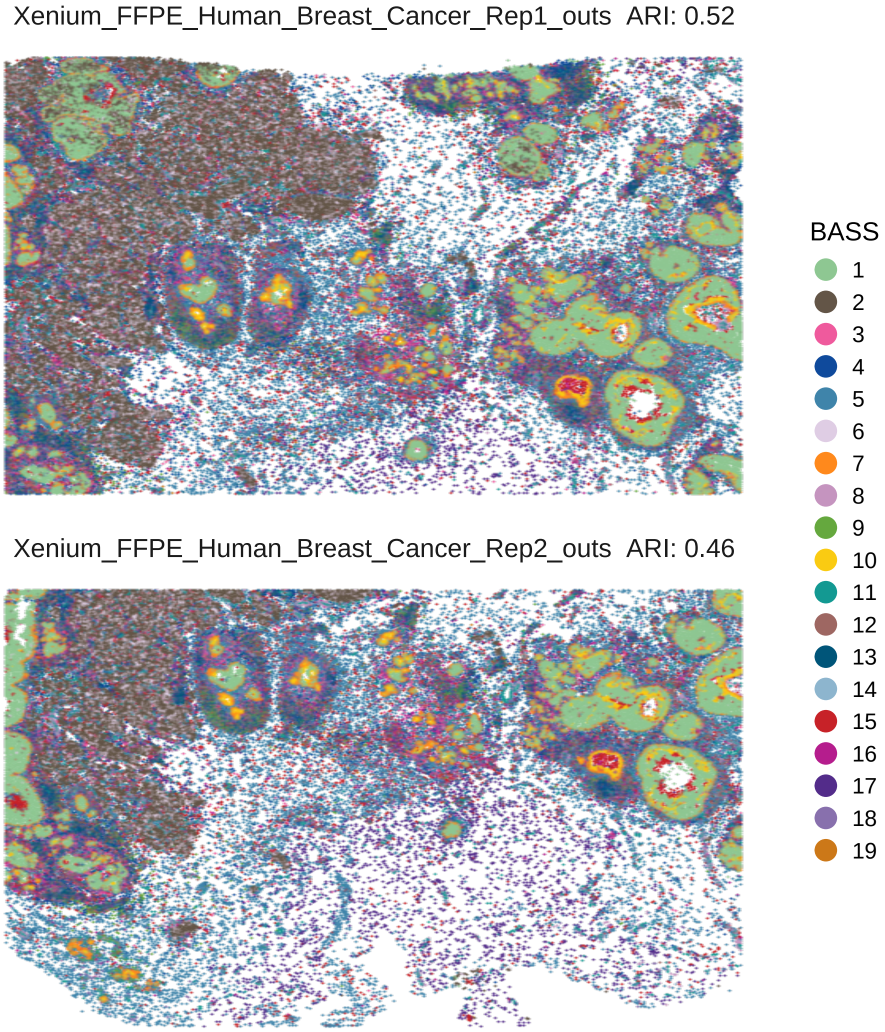


**Suppl Fig. S13:** Spatial plots of BASS clusters in Xenium breast cancer data [6]. ARI between BASS clusters and annotations were calculated for each sample as a measure of clustering concordance.

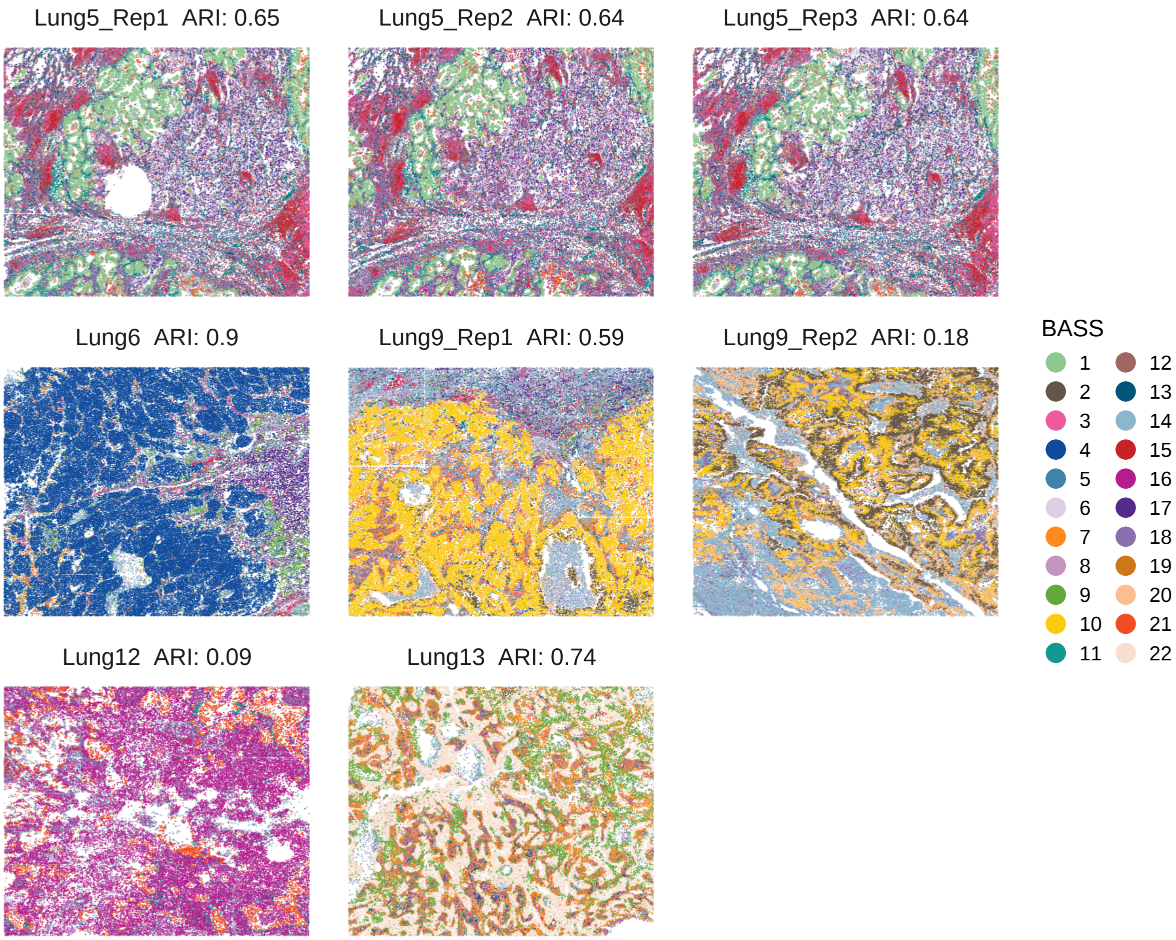


**Suppl Fig. S14:** Spatial plots of BASS clusters in CosMx lung cancer data [5]. ARI between BASS clusters and annotations were calculated for each sample as a measure of clustering concordance.

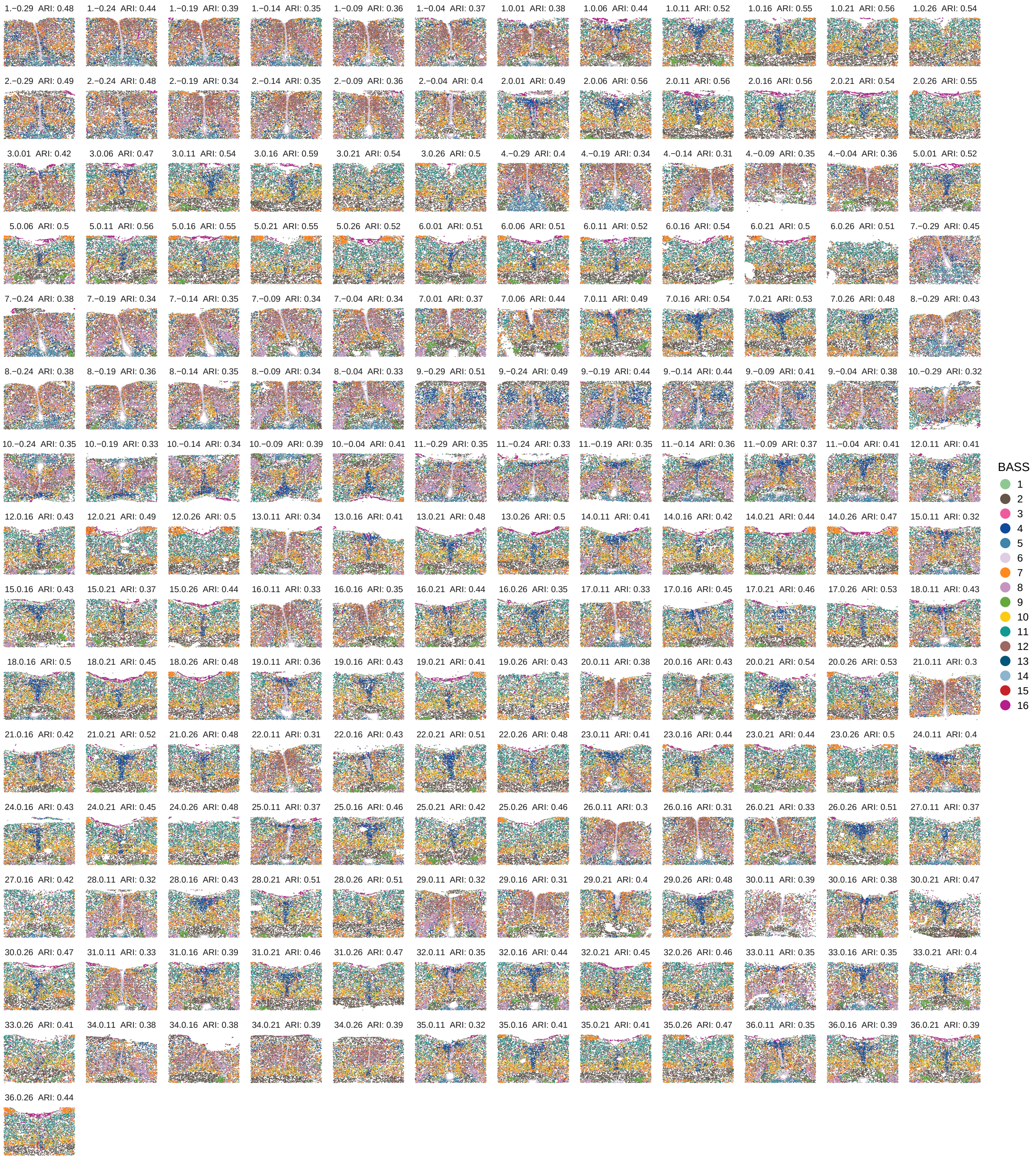


**Suppl Fig. S15:** Spatial plots of BASS clusters in MERFISH mouse hypothalamus preoptic region data [9]. ARI between BASS clusters and annotations were calculated for each sample as a measure of clustering concordance.

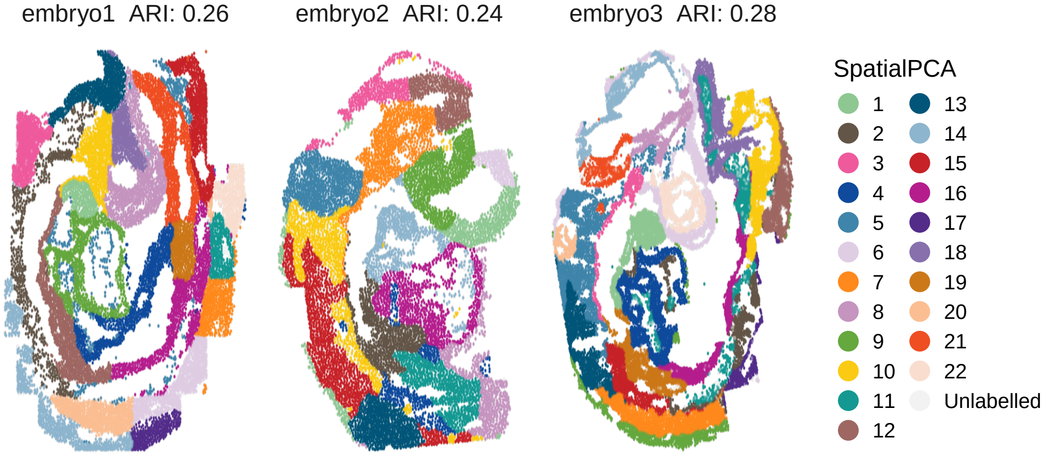


**Suppl Fig. S16:** Spatial plots of SpatialPCA clusters in seqFISH mouse embryo data [1]. ARI between SpatialPCA clusters and annotations were calculated for each sample as a measure of clustering concordance.

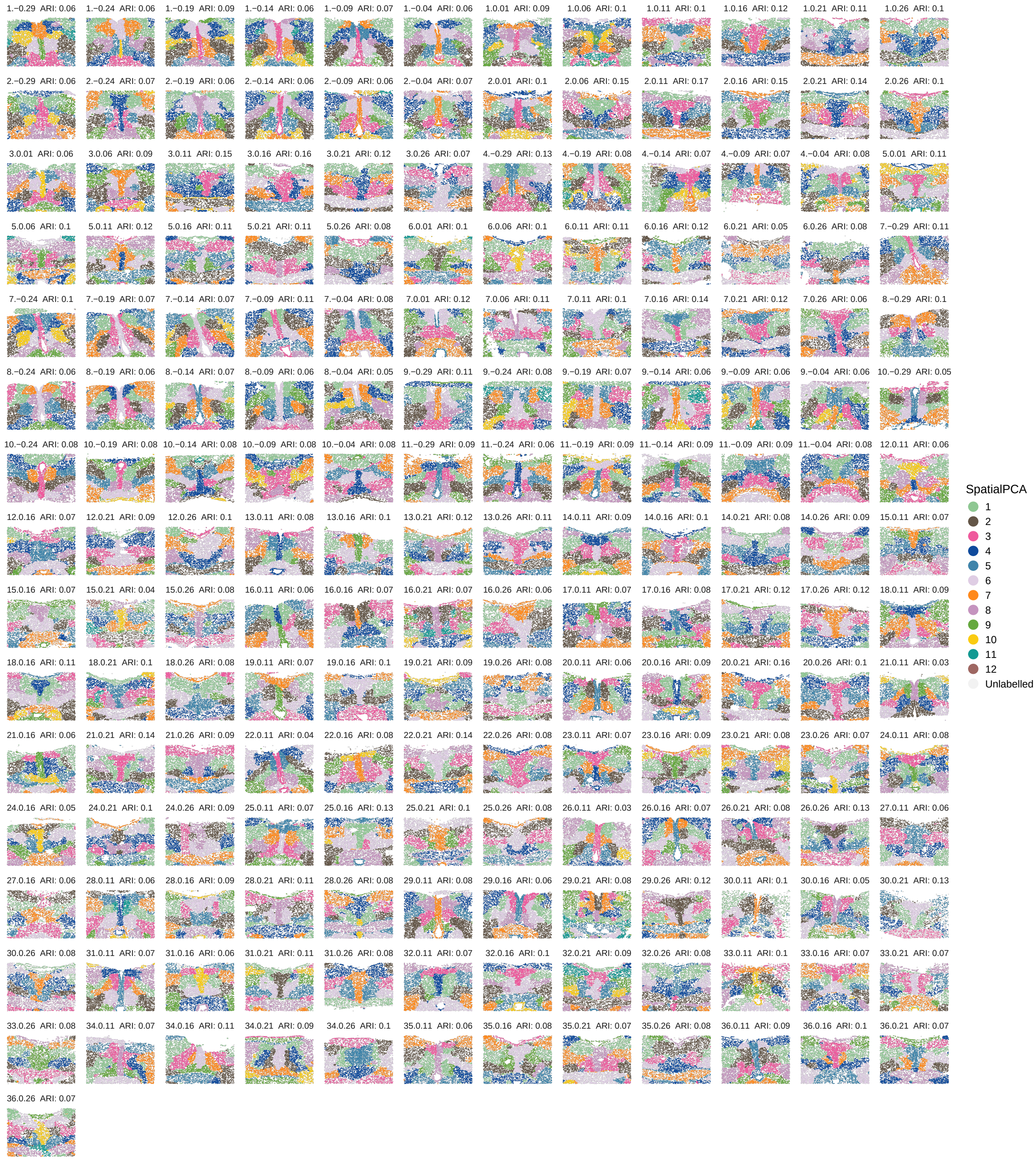


**Suppl Fig. S17:** Spatial plots of SpatialPCA clusters in MERFISH mouse hypothalamus preoptic region data [9]. ARI between SpatialPCA clusters and annotations were calculated for each sample as a measure of clustering concordance.

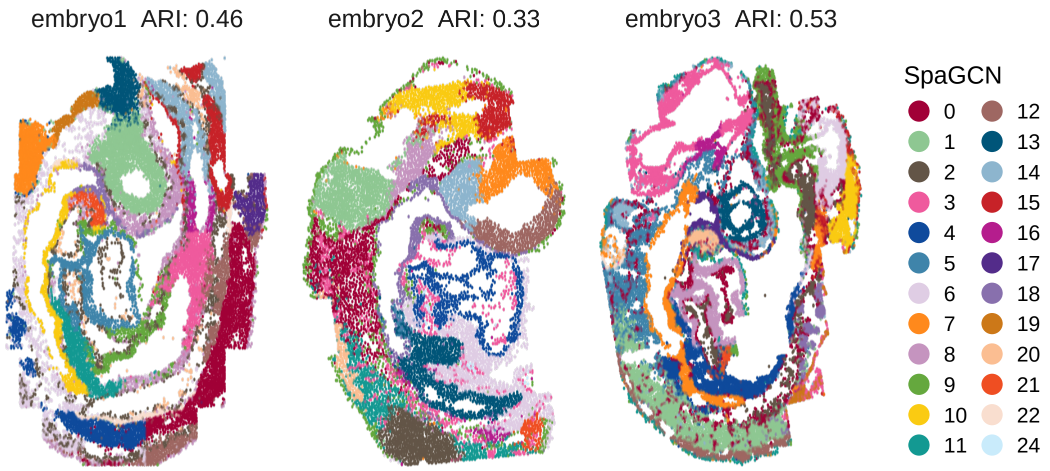


**Suppl Fig. S18:** Spatial plots of SpaGCN clusters in seqFISH mouse embryo data [1]. ARI between SpaGCN clusters and annotations were calculated for each sample as a measure of clustering concordance.

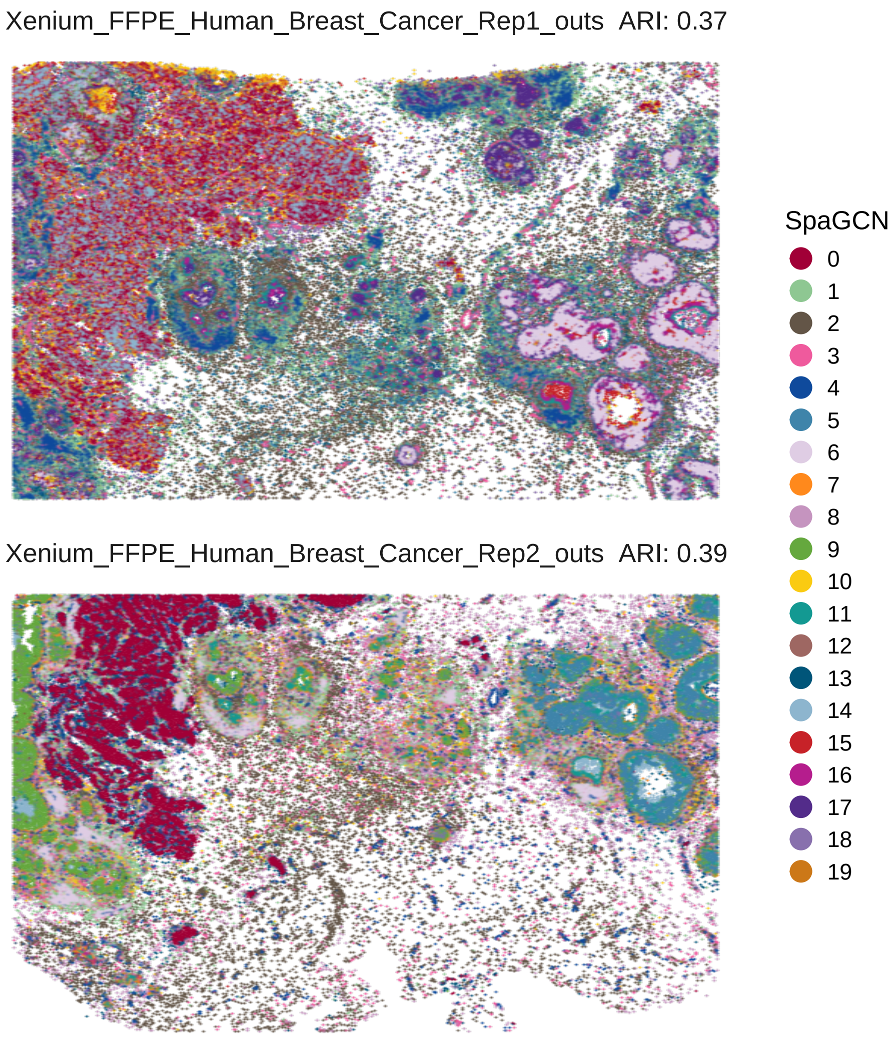


**Suppl Fig. S19:** Spatial plots of SpaGCN clusters in Xenium breast cancer data [6]. ARI between SpaGCN clusters and annotations were calculated for each sample as a measure of clustering concordance.

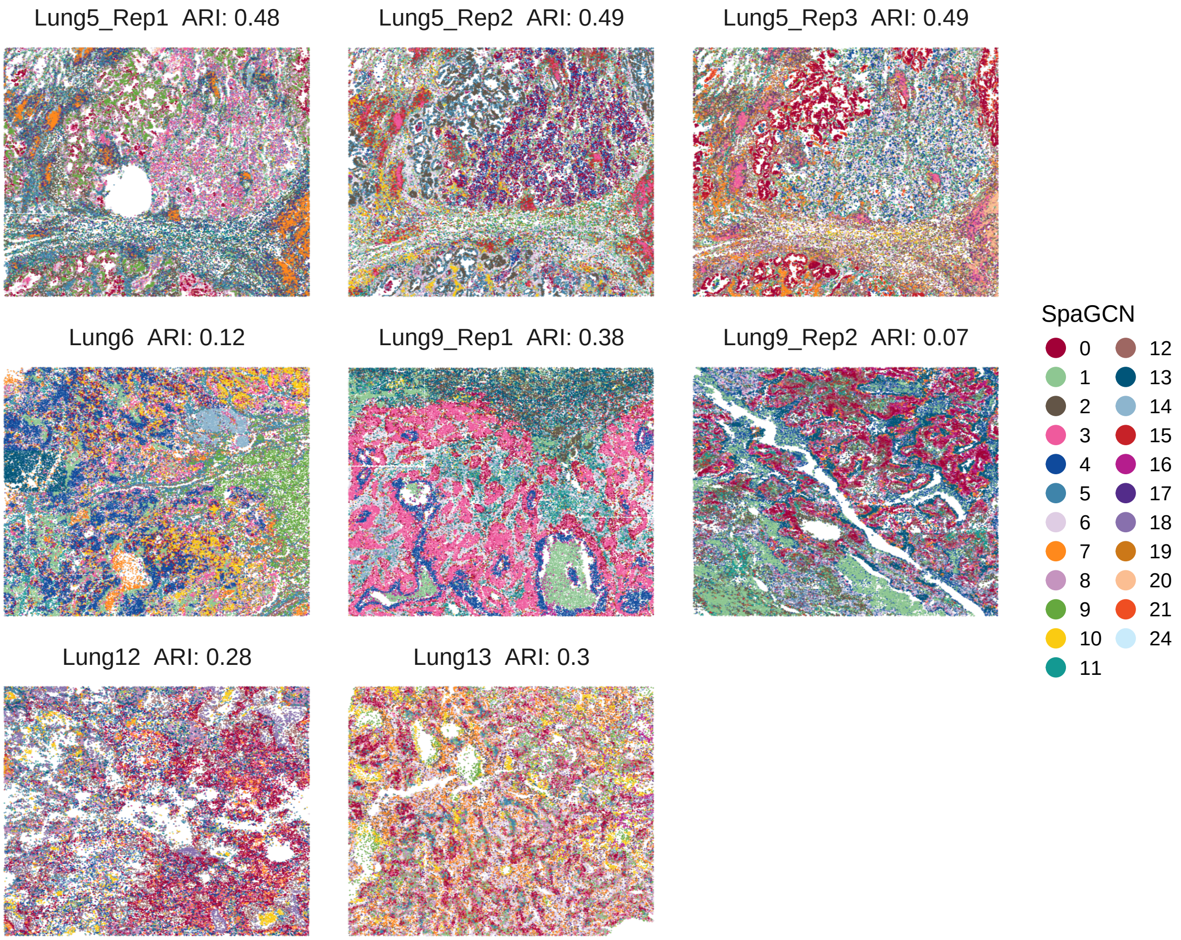


**Suppl Fig. S20:** Spatial plots of SpaGCN clusters in CosMx lung cancer data [5]. ARI between SpaGCN clusters and annotations were calculated for each sample as a measure of clustering concordance.

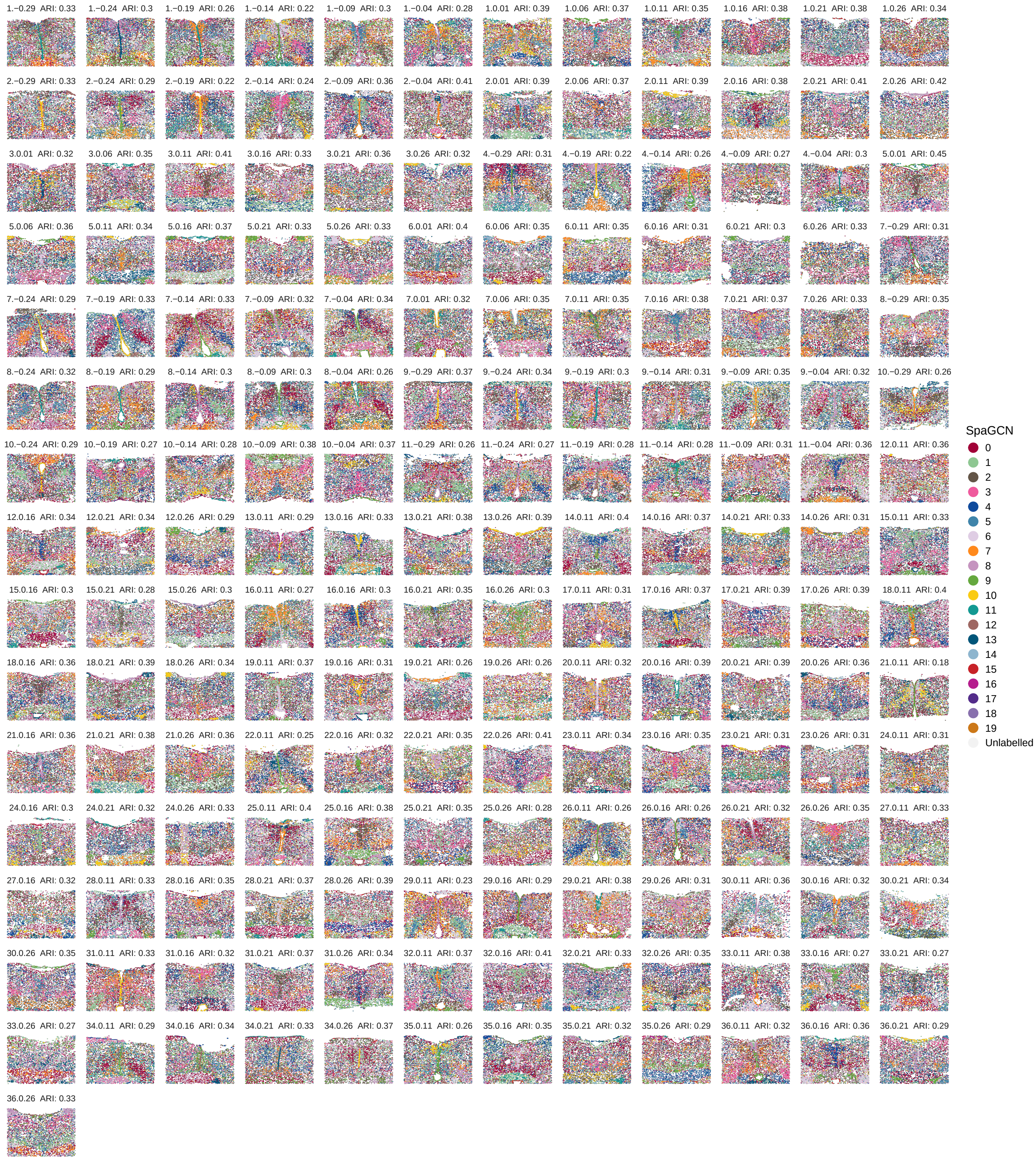


**Suppl Fig. S21:** Spatial plots of SpaGCN clusters in MERFISH mouse hypothalamus preoptic region data [9]. ARI between SpaGCN clusters and annotations were calculated for each sample as a measure of clustering concordance.

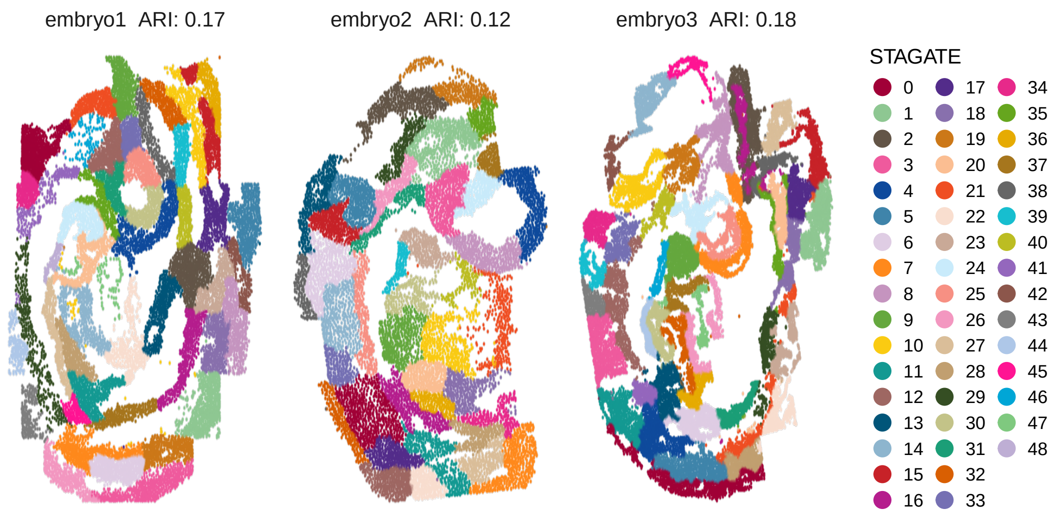


**Suppl Fig. S22:** Spatial plots of STAGATE clusters in seqFISH mouse embryo data [1]. ARI between STAGATE clusters and annotations were calculated for each sample as a measure of clustering concordance.

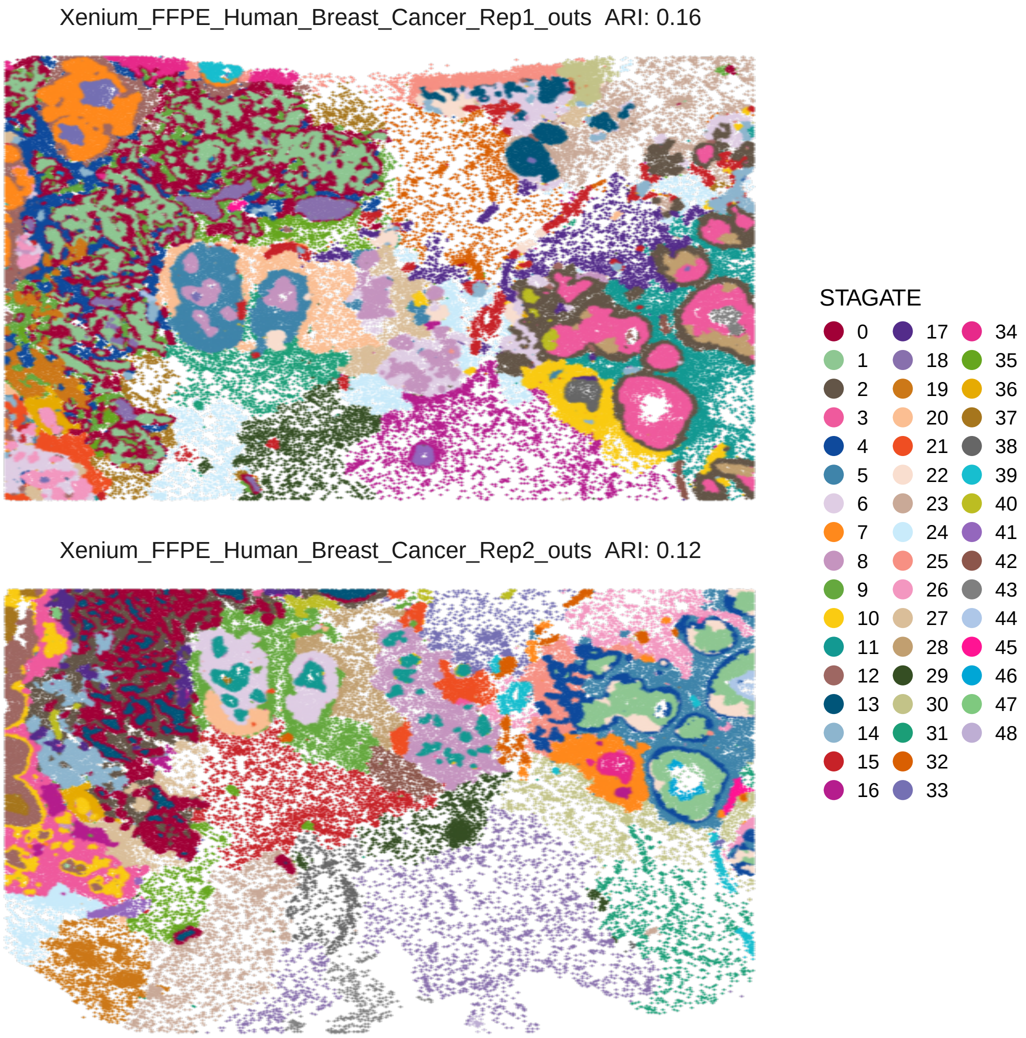


**Suppl Fig. S23:** Spatial plots of STAGATE clusters in Xenium breast cancer data [6]. ARI between STAGATE clusters and annotations were calculated for each sample as a measure of clustering concordance.

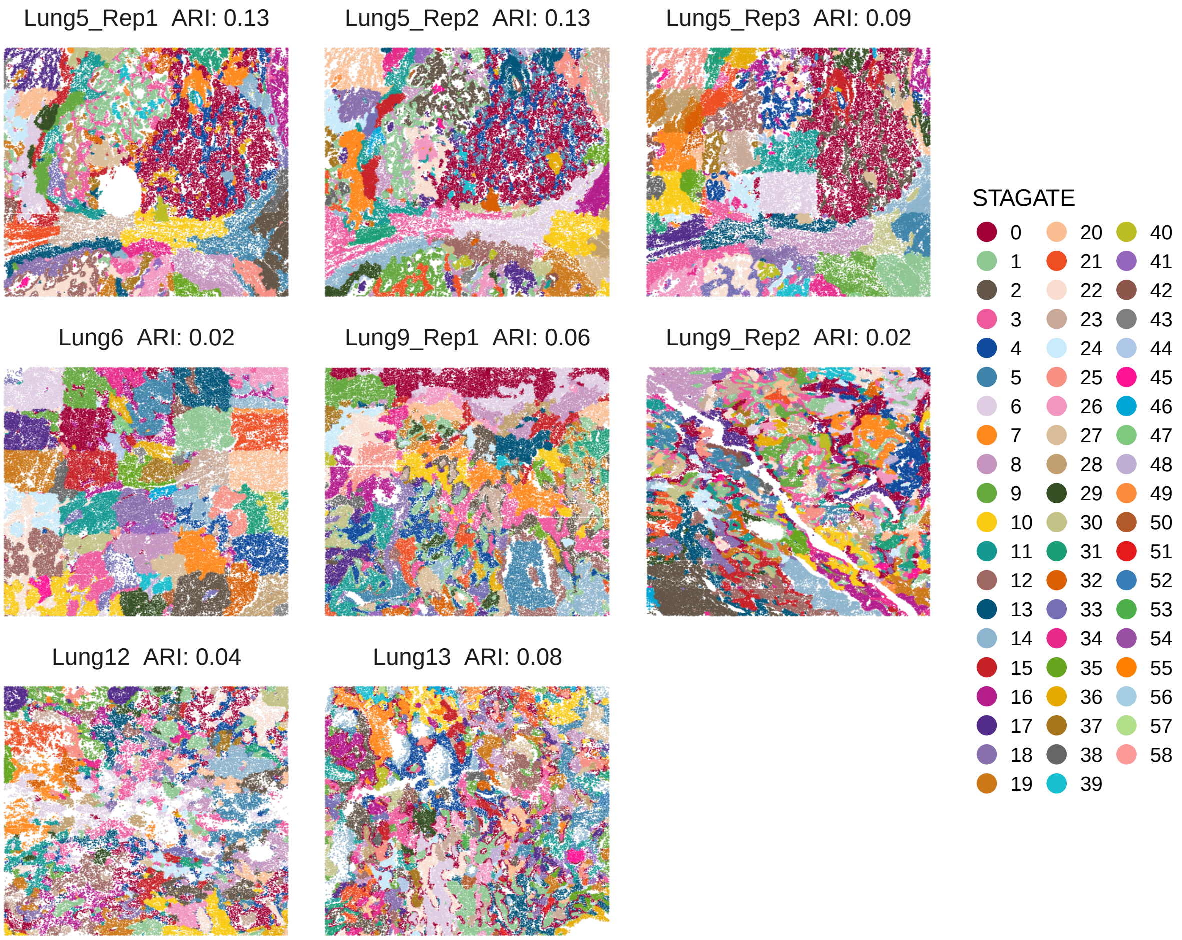


**Suppl Fig. S24:** Spatial plots of STAGATE clusters in CosMx lung cancer data [5]. ARI between STAGATE clusters and annotations were calculated for each sample as a measure of clustering concordance.

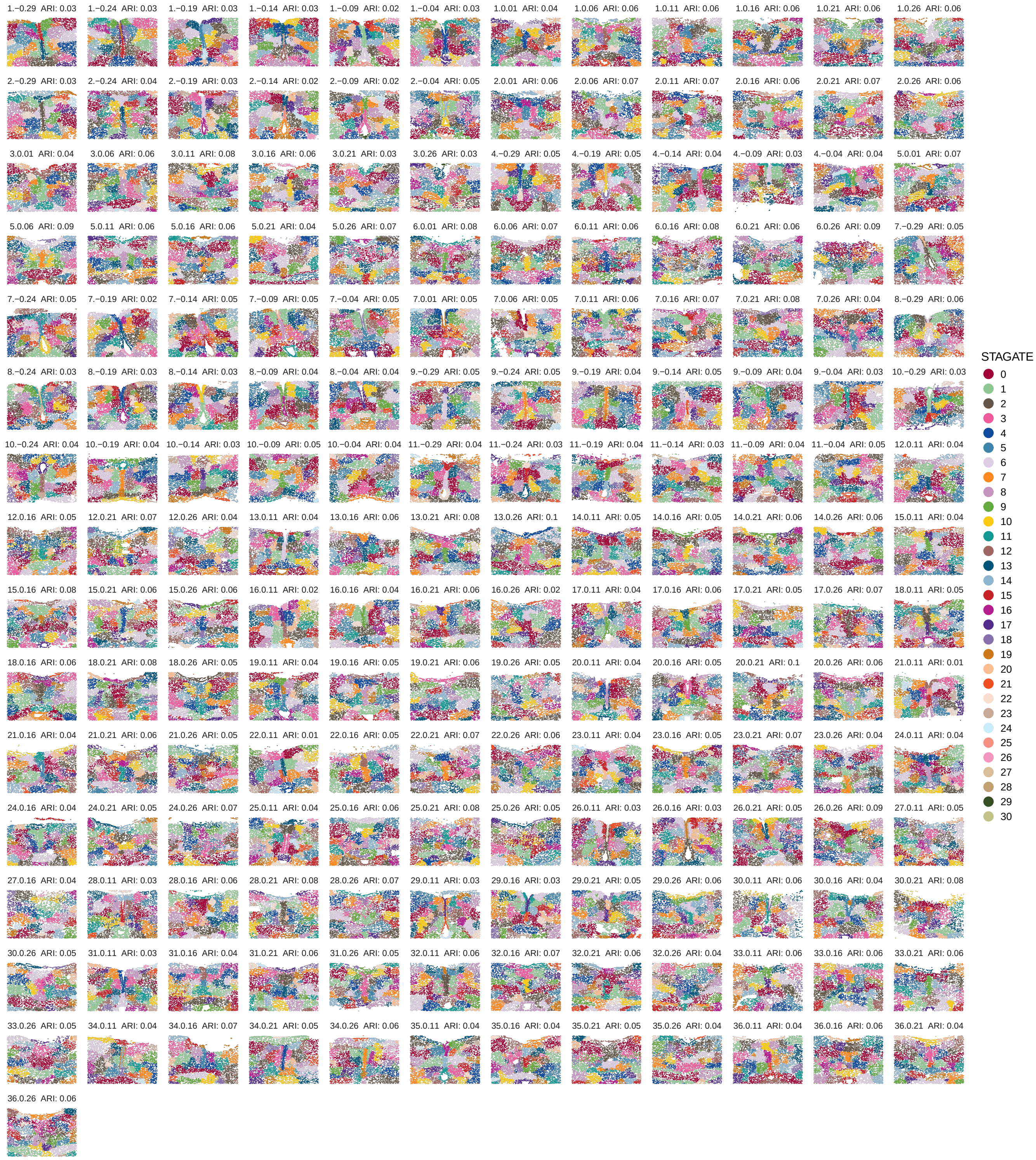


**Suppl Fig. S25:** Spatial plots of STAGATE clusters in MERFISH mouse hypothalamus preoptic region data [9]. ARI between STAGATE clusters and annotations were calculated for each sample as a measure of clustering concordance.

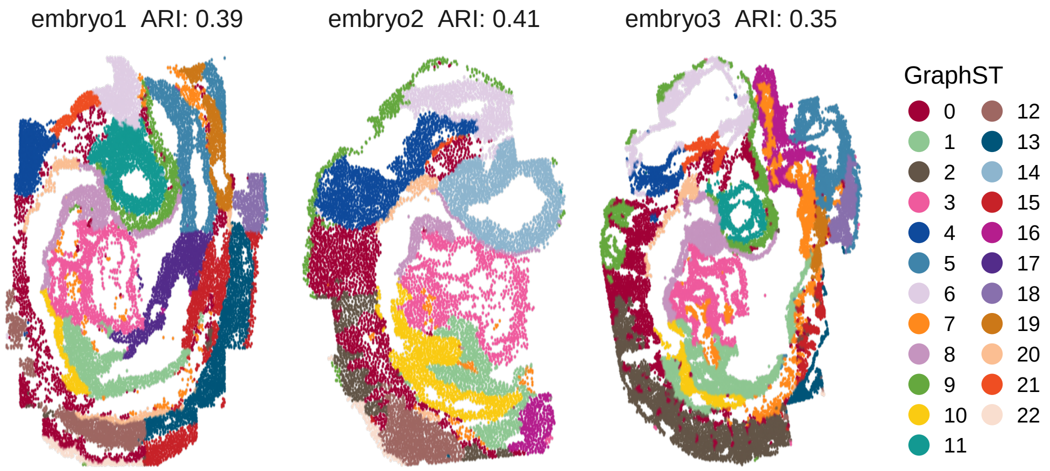


**Suppl Fig. S26:** Spatial plots of GraphST clusters in seqFISH mouse embryo data [1]. ARI between GraphST clusters and annotations were calculated for each sample as a measure of clustering concordance.

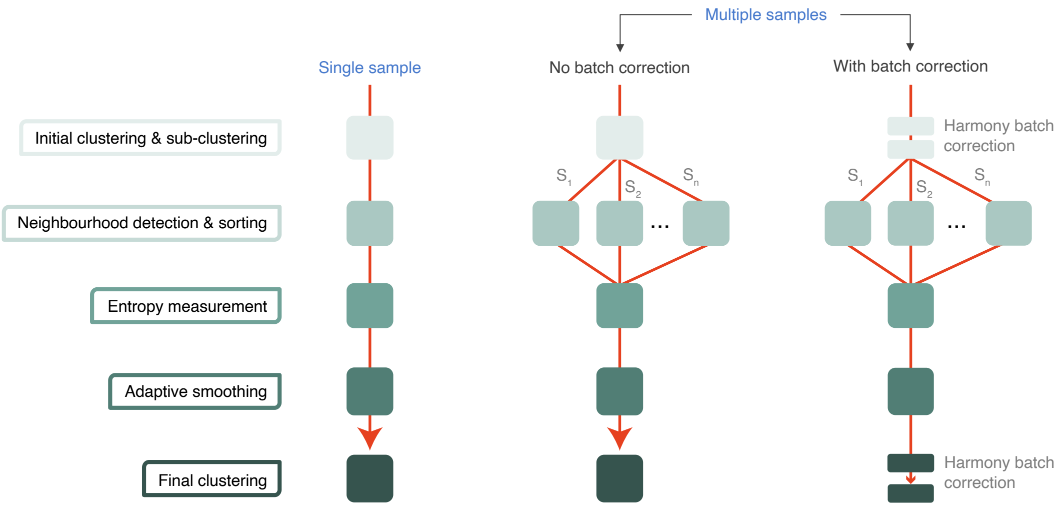


**Suppl Fig. S27:** Schematic overview of ClustSIGNAL's modular algorithm. It involves five sequential steps that differ based on application to single sample analysis, multi-sample analysis with no batch correction, and multi-sample analysis with batch correction.

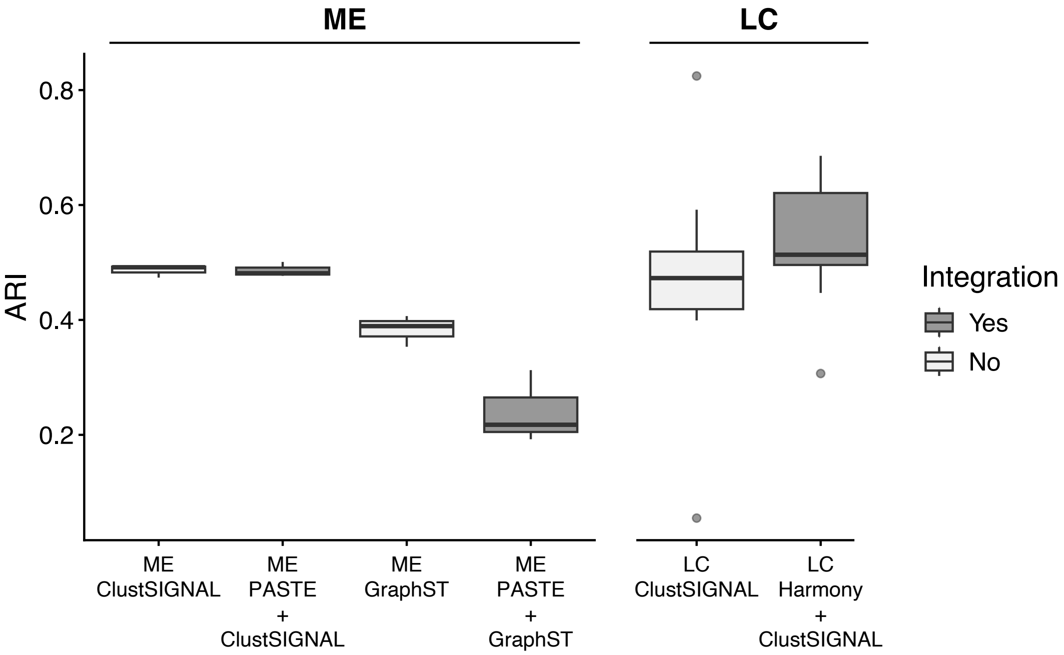


**Suppl Fig. S28:** Box plot showing benchmarking of ClustSIGNAL's multi-sample analysis. Its multi-sample approach was compared with and without PASTE [2] spatial alignment of 3 samples from mouse embryo (ME) data [1] as well as with and without Harmony [4] batch correction of 8 samples from lung cancer (LC) data [5]. ClustSIGNAL was also compared to GraphST, which requires spatial alignment of samples for multi-sample analysis.

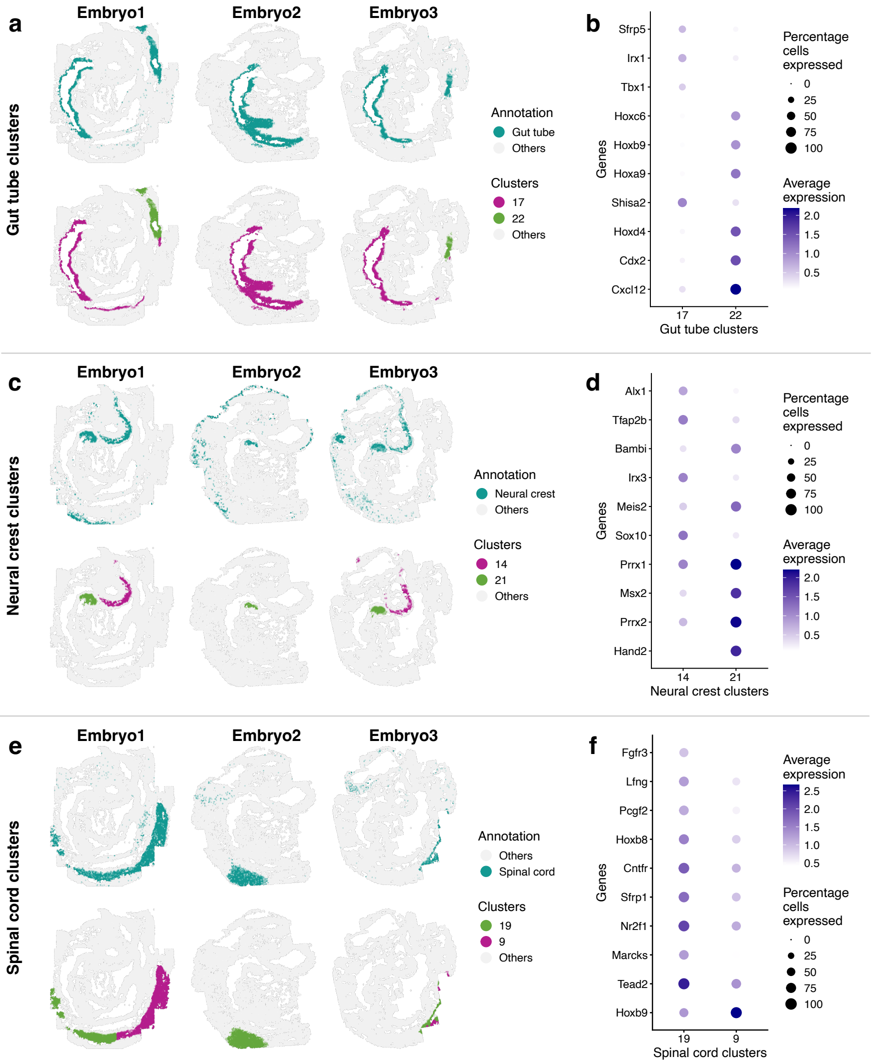


**Suppl Fig. S29:** Spatial plots and dot plots highlighting **(a, b)** gut tube, **(c, d)** neural crest, and **(e, f)** spinal cord regions of mouse embryo (ME) data [1] showing spatial arrangement of prior annotations, region-specific clusters generated by ClustSIGNAL, and top 10 marker genes in each cluster associated with a specific region.

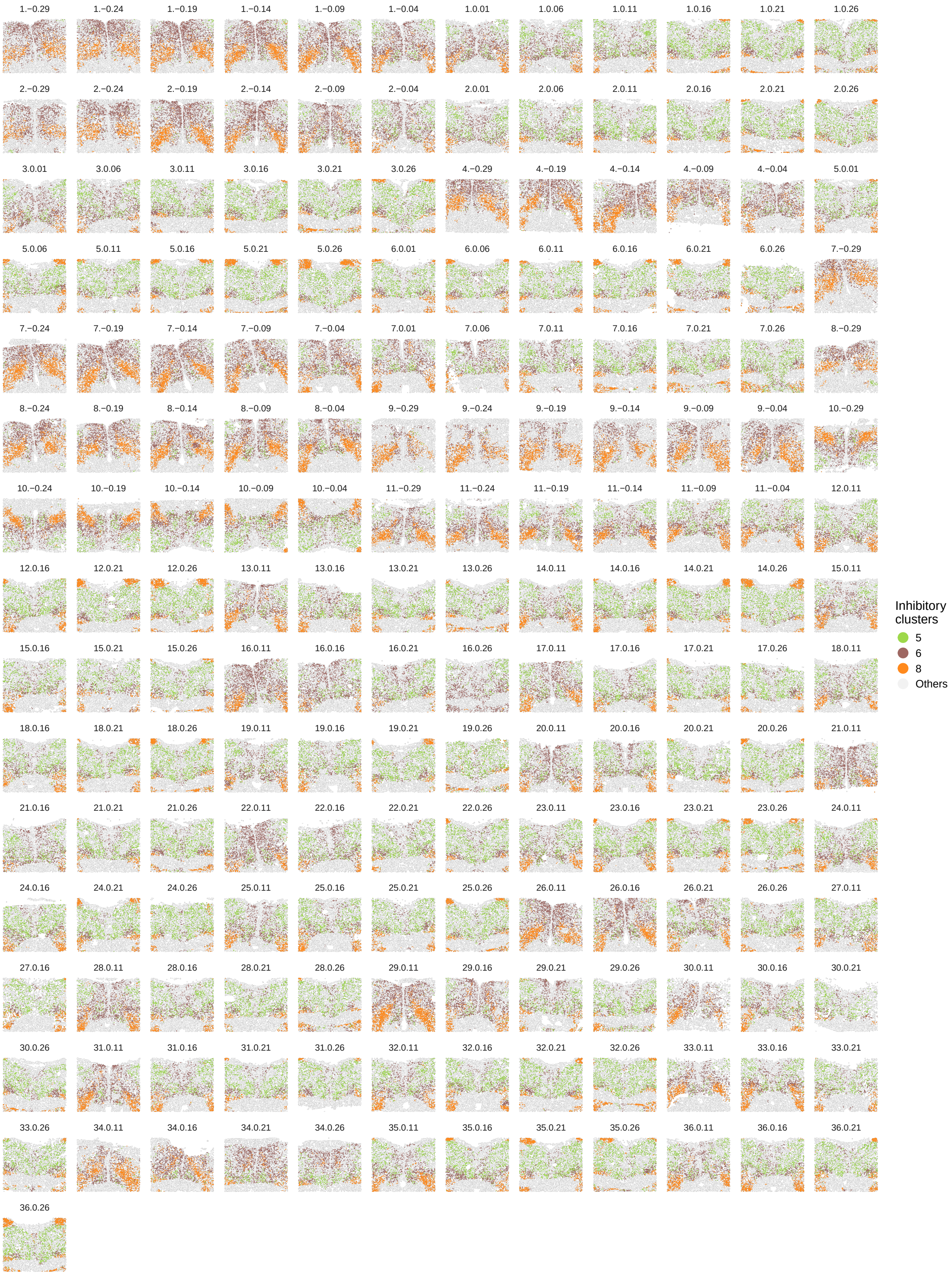


**Suppl Fig. S30:** Spatial plots of Inhibitory neuron clusters identified by ClustSIGNAL in mouse hypothalamus (MH) preoptic region data [9].

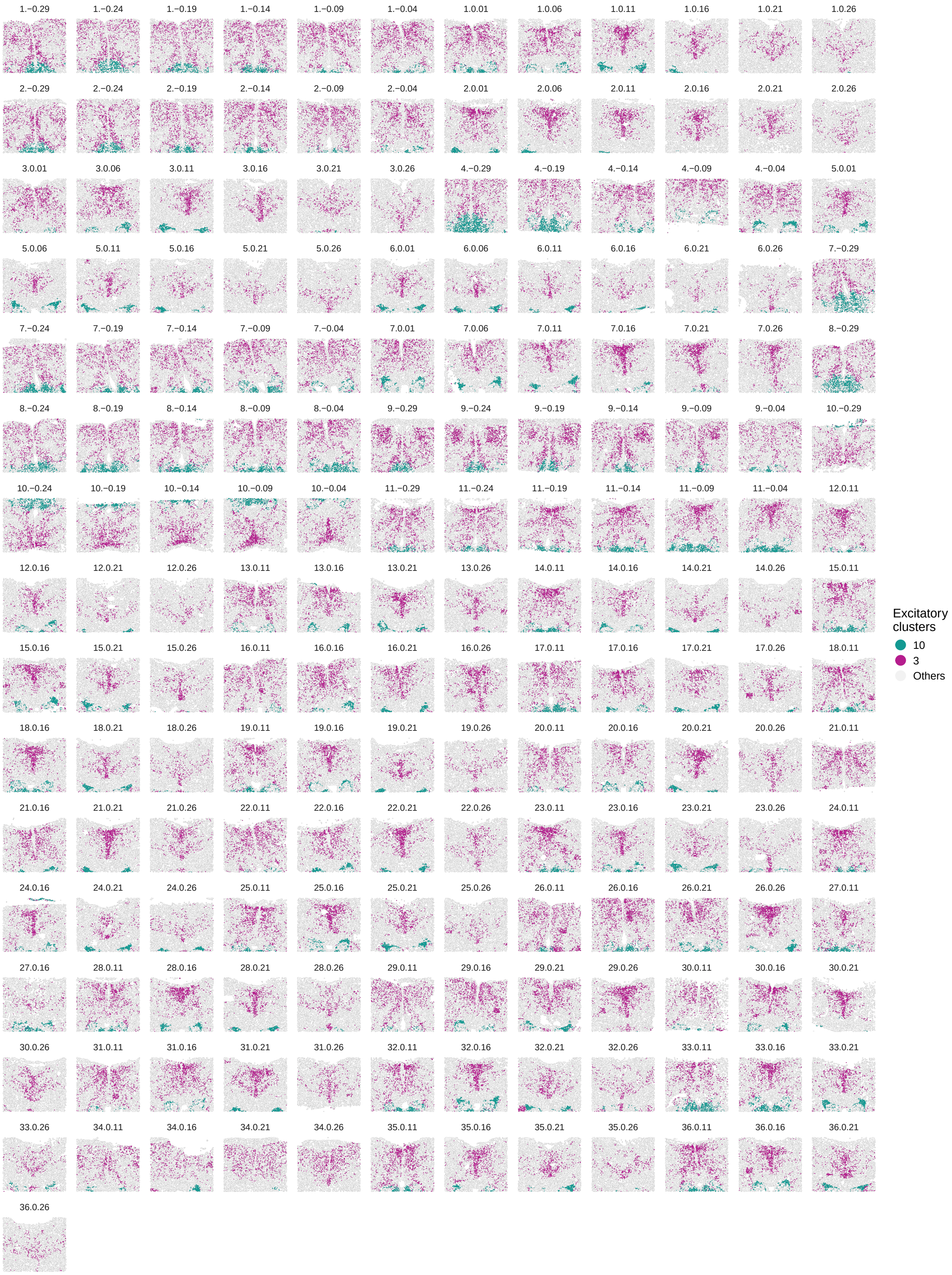


**Suppl Fig. S31:** Spatial plots of Excitatory neuron clusters identified by ClustSIGNAL in mouse hypothalamus (MH) preoptic region data [9].

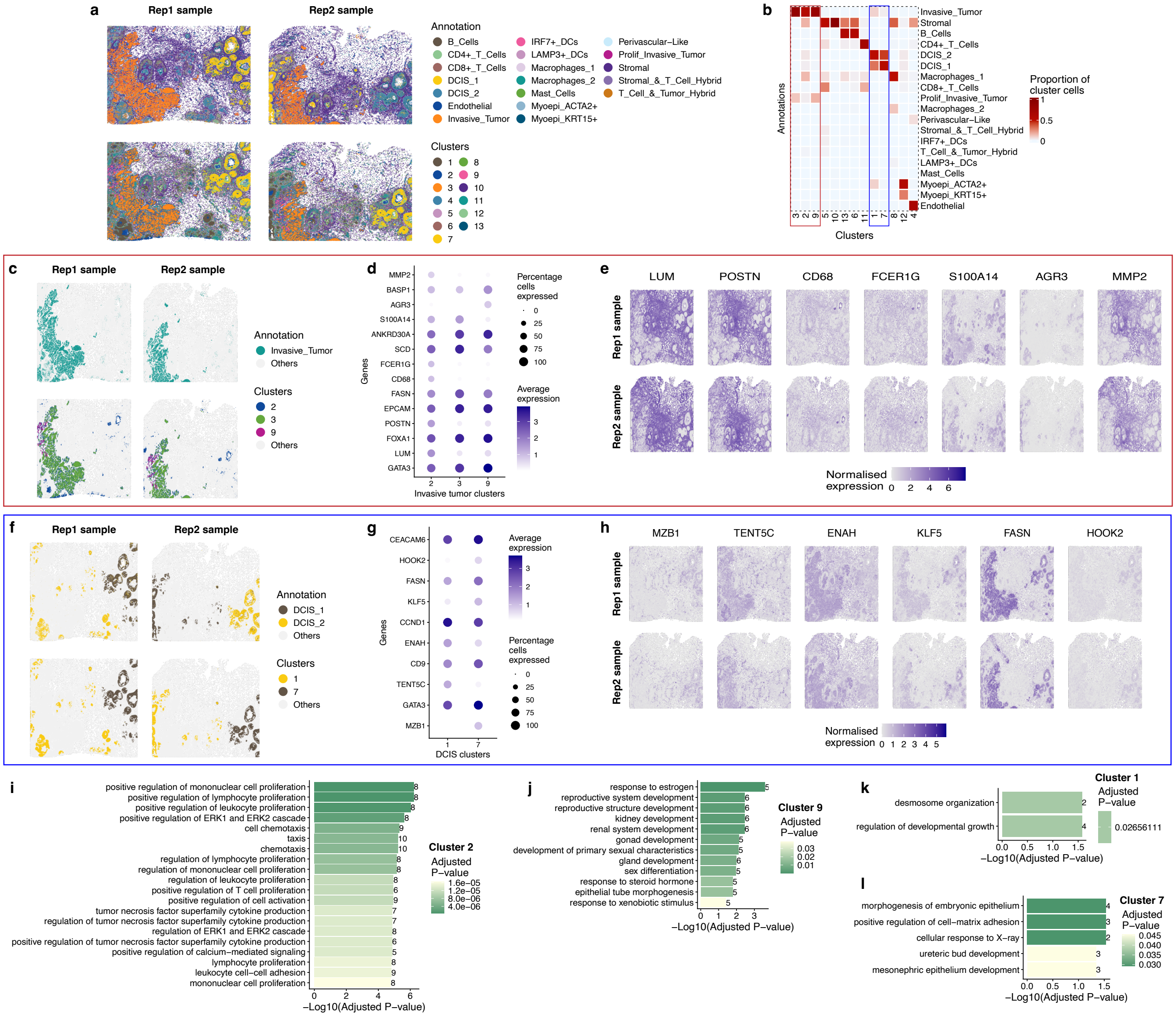

**Suppl Fig. S32:** Analysis of ClustSIGNAL clusters in BR data [6]. **(a)** Spatial plots of prior annotations (top) and ClustSIGNAL clusters (bottom) in the two BR samples. **(b)** Heatmap showing cluster composition, i.e., proportion of cluster cells belonging to each prior annotation. **(c-e)** Analysis of Invasive_Tumor clusters showing **(c)** spatial plots of Invasive_Tumor annotation (top) and related clusters (bottom), **(d)** dot plot of top 10 marker genes in each cluster, and **(e)** spatial plots of distinct marker gene expression. **(f-h)** Analysis of DCIS_1 and DCIS_2 clusters showing **(f)** spatial plots of DCIS_1 and DCIS_2 annotations (top) and related clusters (bottom), **(g)** dot plot of top 10 marker genes in each cluster, and **(h)** spatial plots of distinct marker gene expression. **(i-l)** Pathway enrichment plots showing biological processes (y-axes) associated with the **(i, j)** Invasive_Tumor and **(k, l)** DCIS cluster genes. Only the clusters with significant pathway outputs are shown.

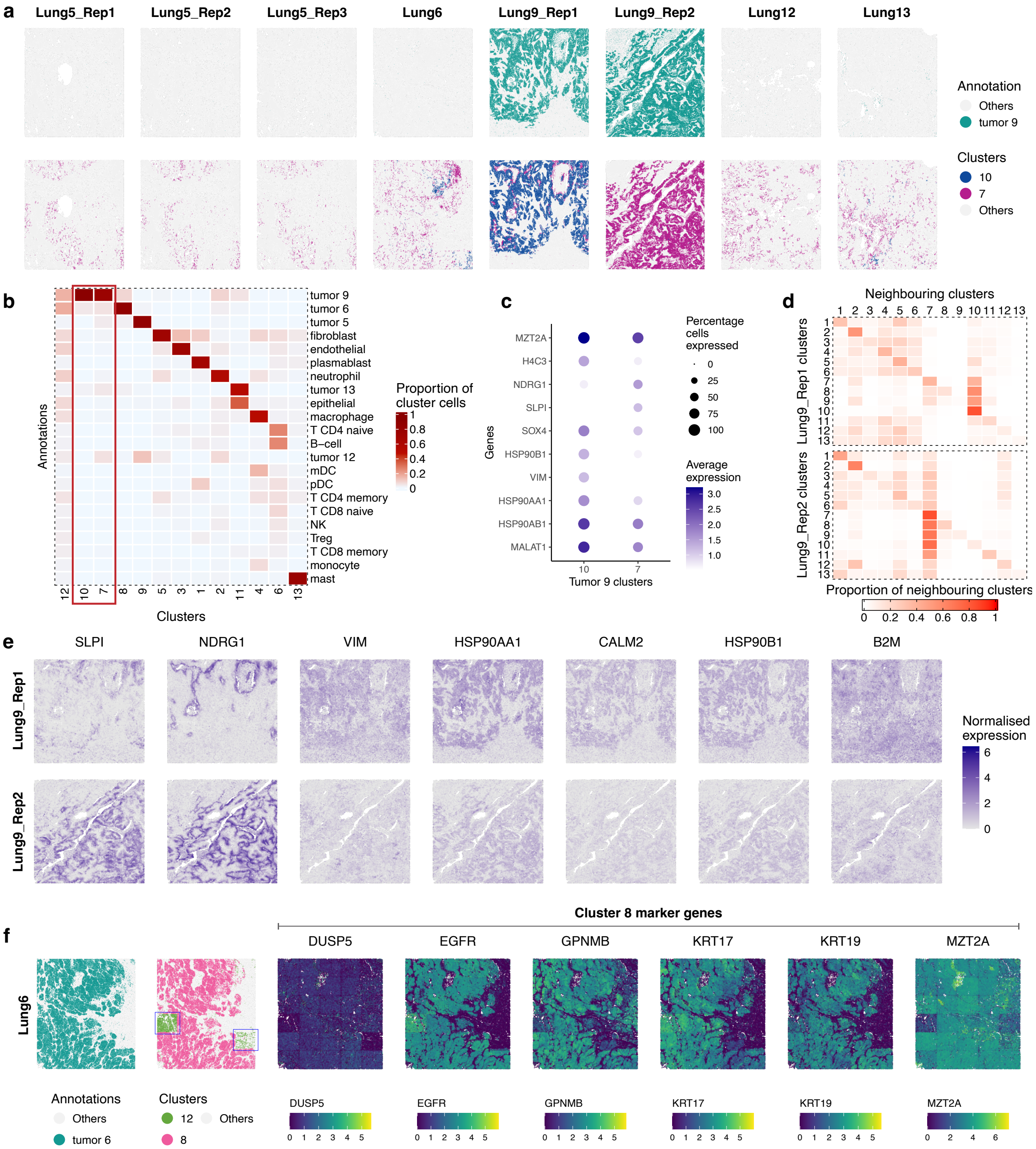


**Suppl Fig. S33:** Analysis of ClustSIGNAL clusters in LC data. **(a)** Spatial plots of tumor 9 annotation (top) and related clusters (10, 7; bottom) in all samples. **(b)** Heatmap showing cluster composition, i.e., proportion of cluster cells associated with each annotation. **(c)** Dot plot of top 10 marker genes in each cluster. **(d)** Co-localisation heatmaps of index clusters (y-axis) and neighbouring clusters (x-axis) in Lung9 samples showing proportion of cells in the neighbourhood of index cluster cells that belonged to specific neighbouring clusters. **(e)** Spatial plots of marker gene expression of tumor 9 clusters **(c)**. **(f)** Spatial plots of tumor 6 cells (first plot), tumor 6-associated clusters (8, 12; second plot), and the marker genes of cluster 8 in Lung6 sample. Cluster 8 was the main tumor 6 cluster in this sample, and field of view batch effects led to partitioning of some tumor 6 cells into a distinct cluster 12.

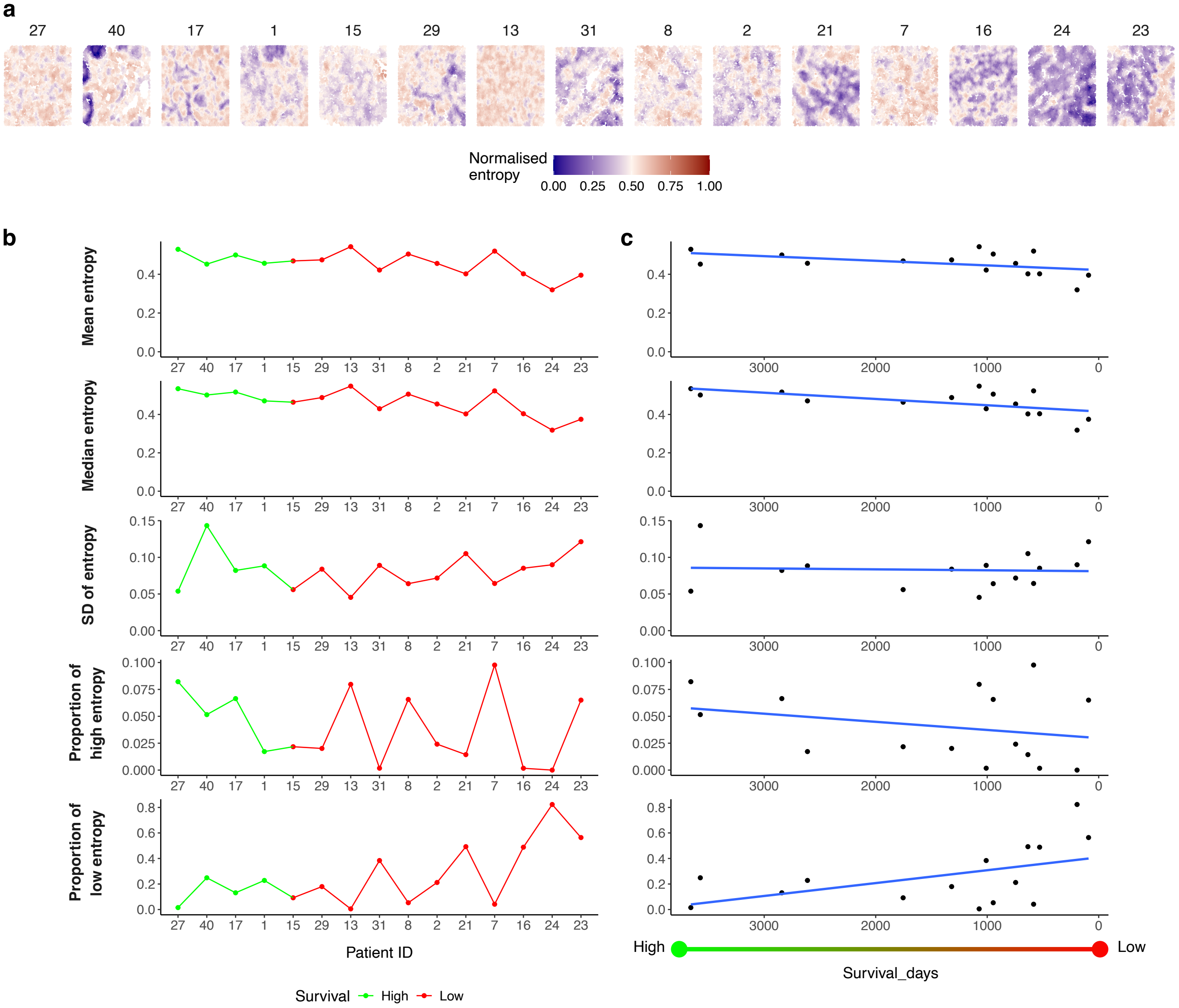


**Suppl Fig. S34:** Analysis of ClustSIGNAL entropy values from a breast cancer spatial proteomics dataset [7]. **(a)** Spatial plots of normalised entropy values of 15 breast cancer samples. **(b)** Plot showing entropy-based metrics (y-axes) plotted for each sample. Samples are arranged and coloured by survival time: Low survival time (< 5 years) in red and High survival time (> 5 years) in green. **(c)** Scatter plot of entropy-based metrics (y-axes) *vs* survival time in days (x-axes), with a linear trend line highlighting the association between the two parameters.
